# Requirement of DNMT1 to orchestrate epigenomic reprogramming during NPM-ALK driven T cell lymphomagenesis

**DOI:** 10.1101/2020.04.09.033373

**Authors:** Elisa Redl, Raheleh Sheibani-Tezerji, Patricia Hamminger, Gerald Timelthaler, Melanie Rosalia Hassler, Sabine Lagger, Thomas Dillinger, Lorena Hofbauer, Maša Zrimšek, Andreas Tiefenbacher, Michael Kothmayer, Charles H. Dietz, Bernard H. Ramsahoye, Lukas Kenner, Christoph Bock, Christian Seiser, Wilfried Ellmeier, Gabriele Schweikert, Gerda Egger

## Abstract

Malignant transformation depends on genetic and epigenetic events that result in a burst of deregulated gene expression and chromatin changes. To dissect the sequence of events in this process, we used a T cell-specific lymphoma model based on the human oncogenic NPM-ALK translocation. We find that transformation of T cells shifts thymic cell populations to an undifferentiated immunophenotype, which occurs only after a period of latency, accompanied by induction of the MYC-NOTCH1 axis and deregulation of key epigenetic enzymes. We discover aberrant DNA methylation patterns, overlapping with regulatory regions, plus a high degree of epigenetic heterogeneity between individual tumors. In addition, ALK positive tumors show a loss of collaborative methylation patterns of neighboring CpG sites. Notably, deletion of the maintenance DNA methyltransferase DNMT1 completely abrogates lymphomagenesis in this model, despite oncogenic signaling through NPM-ALK, suggesting that faithful maintenance of tumor-specific methylation through DNMT1 is essential for sustained proliferation and tumorigenesis.

**STATEMENT OF SIGNIFICANCE:** Epigenetic alterations are causally involved in tumorigenesis. Here we show that induction of a single human oncogene in murine T cells induces specific deregulation of epigenetic enzymes resulting in epigenomic alterations similar to human tumors. Our findings are of broader implication to understand how epigenomic processes are shaped by oncogene induced transformation.

## INTRODUCTION

Individual tumors and tumor types show a high level of heterogeneity regarding their genetic and epigenetic constitution and in the affected signaling pathways. Characteristically altered patterns of DNA methylation, however, are a universal hallmark of human cancer (1): In malignant cells the genome is globally hypomethylated while short CpG dense regions, referred to as CpG Islands (CGI) generally show an increase in methylation (1,2). CGIs are often found in gene promoter regions and hypermethylated CGIs have been associated with the silencing of tumor suppressor genes in diverse cancers. However, linking specific DNA methylation differences with extensive expression changes during tumorigenesis in a cause-and-effect relationship remains challenging, due to the crosstalk of diverse epigenetic regulators. Furthermore, identifying the drivers that target the DNA methylation machinery is equally difficult. Studies in NPM-ALK positive (ALK+) T cell lymphoma, a subgroup of anaplastic large cell lymphoma (ALCL), have implicated the transcription factor STAT3 as a central player in epigenetic regulation (3,4). STAT3 acts by directly and indirectly regulating the expression of the maintenance methyltransferase DNMT1 and by directing all three major methyltransferases (DNMT1, DNMT3A, DNMT3B) to STAT3 binding sites within promoters of genes such as *SHP1* or *IL2RG.* The role of STAT3 as a mediator of DNA methylation of target promoters was supported by recent data, demonstrating a function of acetylated STAT3 for inducing the methylation of tumor suppressor genes in melanoma and breast cancer (5).

On the other hand, it has been recently suggested that disordered methylation patterns in tumors are resulting from stochastic processes and display intra-tumor heterogeneity, which could provide the basis for genetic and epigenetic tumor evolution (6–8).

Interestingly, DNA methylation in tumors is frequently targeted to regions that are associated with H3K27me3 in embryonic stem cells (ESCs), resulting in an epigenetic switch from dynamic Polycomb repressed to more stable DNA methylation based silencing (9–12).

The impact of DNA hypomethylation has been widely studied by using *Dnmt1* hypomorphic alleles, which display reduced DNMT1 protein and activity levels (13–22). Loss of DNA methylation and chromosomal instability seem to promote tumor initiation, whereas hypomethylation of tumor suppressor associated CGIs exerts tumor suppressive effects primarily during tumor progression. Such opposing effects of DNMT1 reduction were observed in different tumor models (19,20). In addition, the *de novo* enzymes DNMT3A and B were shown to be involved in several hematological and solid cancers (23–30). Thus, deregulation or mutation of DNMTs appears to be an essential event in tumorigenesis of various cancers.

In this study, we abrogated tumorigenesis in an NPM-ALK driven T cell lymphoma model by conditionally targeting the DNA methyltransferase *Dnmt1* by *Cd4-Cre* induced deletion. This allowed us to distinguish early NPM-ALK driven events that occur independent of DNA methylation from later transcriptional changes that depend on the methylation machinery. We provide evidence for the cellular events associated with malignant transformation, which follow a period of latency and require the activation of MYC and NOTCH signaling pathways as well as pronounced epigenetic deregulation. Our findings provide further insight how oncogenes drive tumorigenesis by directing large scale transcriptomic and epigenomic alterations. This is an important prerequisite to identify potential therapeutic targets in ALK+ lymphoma and to gain a deeper understanding of large-scale epigenetic rearrangements that drive tumor transformation in general.

## RESULTS

### Induction of ALK dependent tumorigenic pathways following a period of latency

T cell-specific expression of the human oncogenic fusion tyrosine kinase NPM-ALK under the control of the *Cd4* promoter/enhancer element in mice results in 100% transformation and tumor development at a median age of 18 weeks (31). Notably, the time period before tumor onset and age of lethality is highly variable raising important questions about the nature and order of molecular events that need to occur during the latent phase to eventually trigger tumor initiation.

To better understand these steps, we first investigated the molecular state of NPM-ALK induced tumors. We used genome-wide RNA sequencing (RNA-seq) to compare tumor cells in ALK transgenic mice with thymocytes isolated from age-matched wildtype mice. Differential gene expression analysis revealed that ALK tumor cells are characterized by a massive deregulation of gene expression with 2727 genes significantly down- and 1618 genes up-regulated as compared to control (Ctrl) cells, when filtering for differentially expressed genes based on a false discovery rate (FDR)-adjusted p value lower than 0.05 and at least 2-fold changed expression levels (Fig. 1A and Supplemental Table 1).

**Figure 1.**
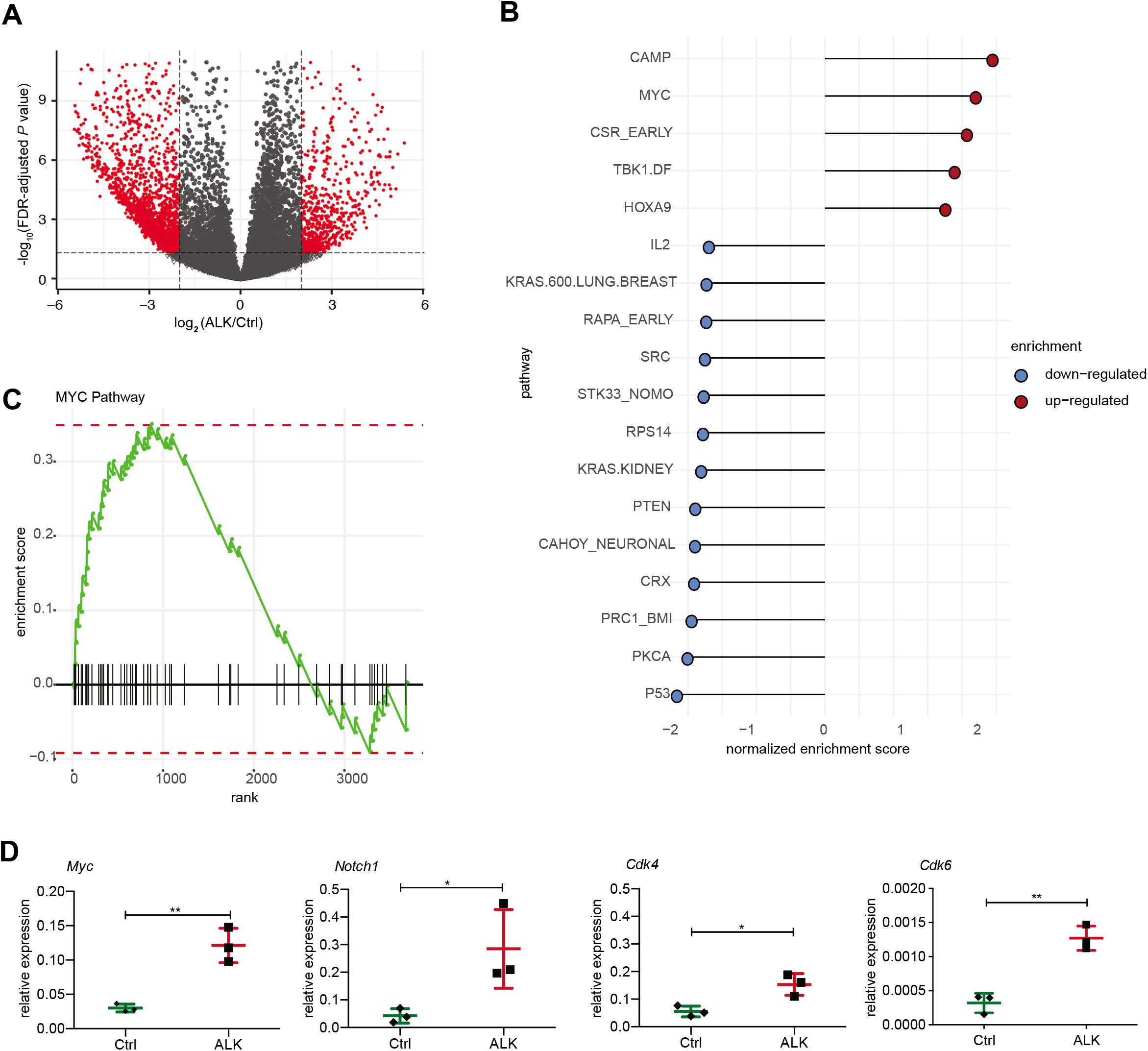
Deregulated gene expression in NPM-ALK tumors. **A**, Volcano plot displaying the differences in gene expression determined by RNA-seq between ALK tumor cells and wild type (Ctrl) thymocytes, where red dots indicate significantly up- and down-regulated genes (permutation test followed by BH correction, FDR<0.05, absolute log2(FC) higher than 1) and grey dots show non-significantly altered genes (not meeting the criteria mentioned above) between these two groups. **B**, Gene set enrichment analysis (GSEA) performed on the significantly up- and down-regulated genes filtered by FDR<0.05 and absolute log_2_(FC)>1 between ALK tumors and Ctrl thymocytes using oncogenic signature gene sets from MSigDB. Pathways associated with downregulated genes are shown in blue, pathways associated with upregulated genes are displayed in red ranked by normalized enrichment score (NES). C, GSEA enrichment of MYC pathway-related genes among significantly deregulated genes filtered by FDR<0.05 and absolute log_2_(FC)>1 between ALK and Ctrl samples. The x-axis shows the differentially expressed genes (DEGs) belonging to the MYC pathway and the y-axis shows positive/negative enrichment scores for up- /down-regulated genes associated with the MYC pathway. D, Analysis of MYC pathway related genes including *Myc, Notch1, Cdk4* and *Cdk6* in Ctrl and ALK tumor samples using RT-qPCR. Analysis was performed in technical and biological triplicates. Data are represented as mean±SD, *p<0.05, ** p<0.01, **** p>0.0001 using unpaired t-test. FC, fold change.

Given the large number of deregulated genes, it is extremely challenging to understand the role and importance of individual expression changes. To identify oncogenic pathways that are associated with ALK dependent tumorigenesis, we first performed gene set enrichment analysis (GSEA) of significantly deregulated genes using oncogenic gene sets from the Molecular Signatures Database (MSigDB) (32,33) (Fig. 1B).

Interestingly, the MYC pathway was among the top upregulated pathways in ALK tumors compared to Ctrl thymocytes (Fig. 1B and 1C), which we also confirmed using quantitative reverse transcription PCR (RT-qPCR) (Fig. 1D). Besides *Myc,* we found the oncogene and Myc-regulator *Notch1* (34) as well as the MYC target genes and cell cycle regulators *Cdk4* and *Cdk6* to be significantly upregulated in ALK tumors. This indicates that MYC signaling is involved in NPM-ALK induced tumorigenesis, as previously observed in human ALK+ ALCL (35).

Furthermore, we found cAMP signaling, which has a role for cell proliferation, differentiation and migration as well as the homeodomain-containing transcription factor HOXA9, which is implicated in hematopoietic stem cell expansion and acute myeloid leukemia (AML), to be upregulated in ALK tumors compared to Ctrl thymocytes (36,37). Additionally, TBK1, an AKT activator and suppressor of programmed cell death was induced in ALK tumors (38).

Among downregulated gene sets, we found genes associated with the tumor suppressors *Atf2, Pten, Pkca, P53* and *Rps14*, a negative regulator of *c-Myc.* Furthermore, genes associated with polycomb repressive complex 1 (PRC1) were also downregulated as well as genes that are upregulated in cells treated with the m-TOR inhibitor Rapamycin (39–44).

### ALK+ tumor cells display an early double-negative (DN) immunophenotype

To get a better understanding of the order of events that lead from NPM-ALK expression to cancer, we next determined the immunophenotype of tumor cells. We used flow cytometry analysis (FACS) to characterize NPM-ALK induced changes in cell composition in transgenic mice compared to control mice at different ages. To specifically analyze ALK+ cells, we combined an intracellular staining for NPM-ALK, with a classical surface staining protocol for common T cell markers. We analyzed thymocytes isolated from thymi of 6- and 18-week old wildtype (Ctrl) and NPM-ALK tumor-free mice (ALK tf), as well as tumor cells from ALK mice that had already undergone transformation (ALK) (Fig. 2A). We found that already in 6-week old transgenic mice almost 100% of T cells were ALK+ while showing no signs of altered thymus morphology. The expression levels of ALK showed a gradual increase from 6 weeks to 18 weeks and were highest in tumor cells compared to untransformed thymocytes (Fig. 2A right panel).

**Figure 2.**
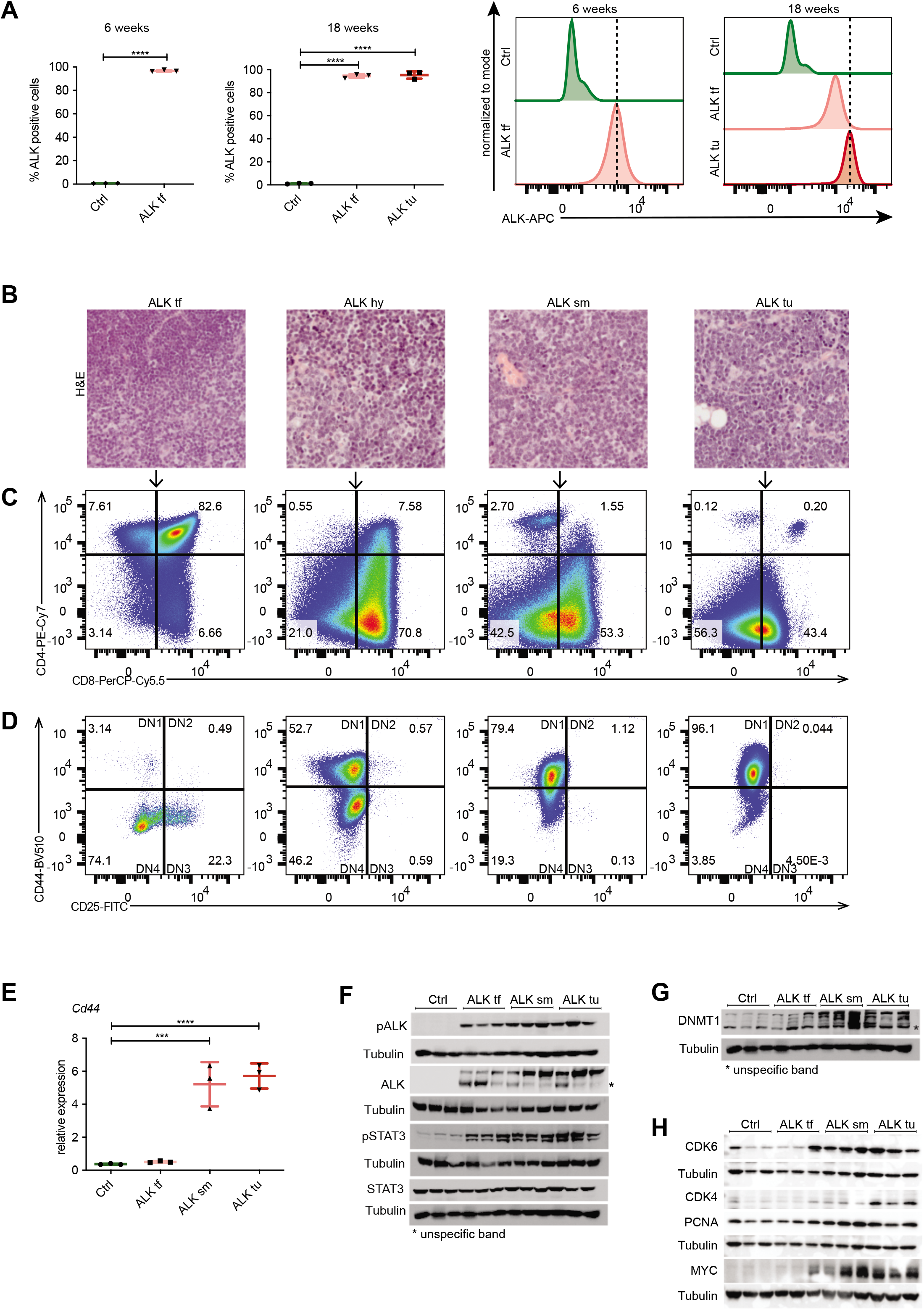
Immunophenotype of ALK tumors. **A**, Intracellular FACS analysis of ALK expression in thymocytes isolated from 6- and 18-week old wild type (Ctrl), NPM-ALK tumor-free (ALK tf) mice compared to ALK+ tumor cells (ALK tu). Quantification of the percentage of ALK+ cells in the three groups (left). Histograms (right) depict ALK expression levels compared to Ctrls. Dotted vertical lines indicate the peaks of ALK expression in ALK+ thymocytes of 6week old ALK mice or ALK+ tumor cells at 18 weeks of age. Data are represented as mean±SD, ****p<0.0001, one-way ANOVA, followed by unpaired t-test, n=3. **B**, Hematoxylin and eosin (HE) stainings of representative thymi of 18-week old NPM-ALK transgenic mice illustrating different stages of ALK tumors including a tumor-free thymus ALK tf; hyperplastic thymus, ALK hy; small tumor, ALK sm; end-stage tumor, ALK tu. **C**, Representative FACS analysis of ALK+ cells isolated from 18-week old tumor-free mice compared to different tumor stages (as in b) gated for CD4 and CD8 expression. **D**, FACS analysis showing the expression of CD44 and CD25 to determine the different double negative (DN) stages of T cell development (DN 1-DN4) in ALK+ cells isolated from 18-week old tumor-free mice and different stages of ALK tumor developing mice (as in b). **E**, RT-qPCR of *Cd44* expression in thymi of 18-week old Ctrl and NPM-ALK tumor-free (ALK tf) transgenic mice as well as early developing tumors (ALK sm) and end-stage tumors (ALK tu) normalized to *Gapdh* expression. Analyses were performed in biological triplicates. Data are represented as mean±SD, ***p<0.001, ****p<0.0001, using one-way ANOVA, followed by unpaired t-test. **F**, pALK, ALK, pSTAT3 and STAT3 protein levels in biological triplicates of thymi of 18-week old Ctrl and NPM-ALK tumor-free mice as well as early and endstage tumors were analyzed by Western blot analysis. Tubulin served as loading control. Asterisks indicate unspecific protein bands. **G**, DNMT1 protein levels in biological triplicates of thymi of 18-week old Ctrl and tumor-free NPM-ALK transgenic mice as well as early and end-stage tumors (as in f) were analyzed by Western blot analysis. Tubulin served as loading control. Asterisks indicate unspecific protein bands. **H**, Protein levels of the cell cycle associated genes CDK4, CDK6 and PCNA as well as the oncogene c-MYC in biological triplicates of thymi of 18-week old Ctrl and NPM-ALK tumor-free mice as well as early and end-stage tumors were examined using Western blot analysis. Tubulin served as loading control.

Despite early expression of ALK, the distribution of T cell subsets was normal in 18-week old ALK tumor-free mice as compared to Ctrls (Supplementary Fig. 1A). In NPM-ALK tumors of 18-week old mice however, we observed the previously reported switch to CD4–CD8+ single-positive (SP) or CD4–CD8–double-negative (DN) subsets (31). Furthermore, a similar fraction of Ctrls and ALK tumor-free thymocytes expressed the T cell receptor beta (TCRβ), while the TCRβ+ fraction was absent in cells isolated from ALK tumors (Supplementary Fig. 1B). These two findings suggest that during the initial latent phase, T cell development progresses normally despite NPM-ALK expression and that the induction of T cell transformation happens thereafter.

ALK+ T cells were also present in spleens from 18-week old Ctrl and ALK tumor-free mice as well as ALK tumor mice (Supplementary Fig. 2A). Before tumor onset, these ALK+ cells were either CD4+ or CD8+ T cells (i.e. TCRβ+), whereas in tumor-bearing mice additional CD4–CD8–TCRβ-negative tumor cells were detectable, suggesting that a fraction of ALK transformed DN tumor cells was able to leave the thymus, or that the TCR expression is silenced after exit from the thymus as suggested previously (45) (Supplementary Fig. 2B). These data suggest that ALK induced transformation lymphomagenesis involves the induction of additional tumorigenic pathways and the repression of tumor suppressive genes.

### Tumor transformation and growth is accompanied by the occurrence of tumor cells with an immature T cell profile

To further investigate the transformation process of thymocytes in NPM-ALK mice, we collected a series of thymi from NPM-ALK transgenic mice at 18 weeks of age, which displayed different sizes and morphology correlating with different tumor stages as determined by histological analysis (Fig. 2B). Immunophenotyping of these thymic samples revealed a gradual switch from CD4+CD8+ DP thymocytes to immature single positive (ISP) thymocytes and DN subsets, correlating with tumor progression (Fig. 2C). Further analysis of the DN population based on CD25 and CD44 expression revealed that in tumor-free thymi, the majority of ALK+ cells were DN3 (CD25+CD44-) and DN4 (CD25–CD44-) stage thymocytes, which correlates with the developmental stages at which *Cd4* driven NPM-ALK expression was induced (Fig. 2D). During tumorigenesis, there was a shift towards the DN 1-like (CD25-CD44+) stage, which was also confirmed by RT-qPCR based on *Cd44* expression (Fig. 2E). Together, these data suggest that ALK tumors are developing from a subpopulation of immature T cells or through reprogramming of DP T cells towards the DN stage.

### DNMT1 and STAT3 are activated during the latent phase

Next we sought to identify events that occur in the latent phase prior to tumor initiation, which are potentially responsible for triggering the observed large-scale downstream reprogramming events. As already demonstrated by flow cytometry analysis, NPM-ALK and its active phosphorylated form pALK were detected by Western blot analysis independent of tumor presence, but with increasing protein levels correlating with tumor progression (Fig. 2F, Supplementary Fig. 3). Global STAT3 levels were unaffected by ALK expression, however, its activated form pSTAT3 was significantly increased upon ALK induction independent of tumor presence, suggesting an early role in the latent phase before tumor onset. Previous work including our own has implicated epigenetic mechanisms in NPM-ALK mediated lymphomagenesis in human cell lines and tumors in part through a direct effect of NPM-ALK and STAT3 signaling for the regulation and targeting of the major DNA methyltransferase DNMT1 (4,46–50). Thus, we investigated DNMT1 protein levels before tumor onset and in different tumor stages from tumor-free to end-term tumors in NPM-ALK transgenic mice in comparison to 18-week old wildtypes. Western blot analysis revealed a slight DNMT1 upregulation already in tumor free mice, which further increased during tumor progression (Fig. 2G). Upregulation of DNMT1 has been closely linked to the cell cycle and cell proliferation (51). To test whether the observed upregulation of DNMT1 expression in ALK tumor samples is associated with deregulation of cell cycle genes, we performed Western blot analysis using antibodies against CDK4 and CDK6 as well as c-MYC and the proliferation marker PCNA (Fig. 2H). All cell cycle related proteins followed the previously detected DNMT1 expression pattern with a very prominent upregulation of CDK6 and c-MYC upon tumor onset in thymi harboring small tumors, which was maintained in end-stage tumors. Together, these results suggest that ALK signaling is already active in pre-tumor stages and leads to an early activation of pSTAT3 and elevated expression of DNMT1 culminating in ALK-dependent transformation in NPM-ALK transgenic mice, which is accompanied by upregulation of cell cycle genes and c-MYC induction.

### Deletion of *Dnmt1* abrogates NPM-ALK dependent tumorigenesis

To investigate the functional role of DNMT1 for ALK driven tumorigenesis in more detail, we intercrossed the *Cd4*-NPM-ALK transgenic mice (ALK) with mice carrying a T cell-specific loss of *Dnmt1* (*Cd4*-Cre)(KO) (52). The resulting strain expresses the human *NPM-ALK* transgene but lacks a functional *Dnmt1* gene in T cells (ALKKO) (Fig. 3A). It was shown previously that deletion of *Dnmt1* via the *Cd4* promoter in the double positive stage of T cell development does not interfere with T cell development (52), thus providing a suitable model to study the function of DNMT1 for ALK dependent transformation of thymocytes. Strikingly, deletion of *Dnmt1* in this model completely abrogated NPM-ALK driven lymphomagenesis as shown by Kaplan Meier survival statistics (Fig. 3B). The life span of ALKKO mice was identical to Ctrl and *Dnmt1* knockout mice, and no aberrant phenotype was detected in ALKKO thymi (Fig. 3C). Effective deletion of *Dnmt1* in KO and ALKKO mice was confirmed on protein level by immunohistochemistry (IHC) and Western blot analysis (Fig. 3D,E). *Dnmt1* knockout in the context of ALK expression did not lead to changes in the relative percentages of DN, DP and CD4SP and CD8SP thymocyte subsets (Supplementary Fig. 4). Notably, abrogation of DNMT1 did not interfere with ALK levels or activity in ALKKO transgenic mice as indicated by similar pALK levels in ALK tumor cells and ALKKO thymocytes nor did it change pSTAT3 levels (Fig. 3E). Thus, our data suggest that depletion of DNMT1 interrupts the chain of events that leads from ALK activation to T cell transformation.

**Figure 3.**
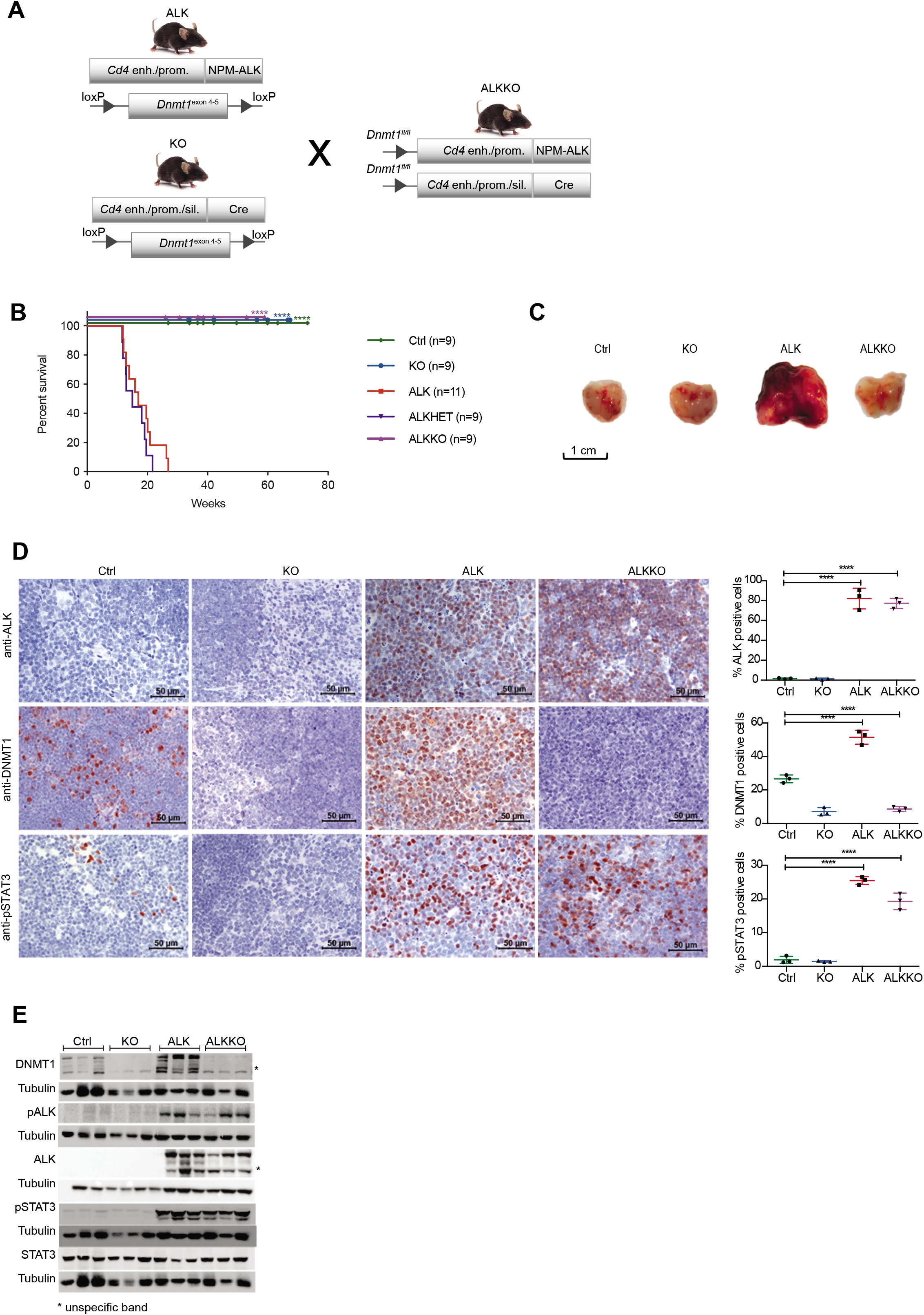
Deletion of *Dnmt1* abrogates lymphomagenesis in NPM-ALK transgenic mice. **A**, Generation of mice with T cell-specific *Cd4*-NPM-ALK expression (ALK) and T cell-specific deletion of *Dnmt1* (KO) or both (ALKKO). *Cd4* enh./prom., *Cd4* enhancer and promoter. *Cd4* enh./prom./sil., *Cd4* enhancer, promoter and silencer. **B**, Kaplan-Meier survival statistics depicting overall survival of *Cd4*-NPM-ALK (ALK), *Cd4*-NPM-ALK *Cd4*-Cre *Dnmt1flox/+* (ALKHET), *Cd4*-NPM-ALK *Cd4*-Cre *Dnmt1flox/flox* (ALKKO) and *Dnmt1loxP/loxP* control mice (Ctrl). ****p<0.0001, Log-rank (Mantel Cox) test, pairwise comparison to ALK. **C**, Morphology of 18-week old Ctrl, KO and ALKKO thymi in comparison to ALK tumors. Pictures were taken immediately after organ collection. **D**, Protein expression of NPM-ALK, DNMT1 and pSTAT3 was analyzed by immunohistochemistry staining in Ctrl, KO, ALKKO thymi as well as in ALK tumors. Pictures are representatives of biological triplicates. Graphs below the images depict quantification of stainings using Definiens TissueStudio 4.2 software. Data are represented as mean±SD, **p<0.01, ***p<0.001 and ****p<0.0001, using one-way ANOVA, followed by unpaired t-test. **E**, DNMT1, ALK, pALK, STAT3 and pSTAT3 protein levels in thymi of 18-week old Ctrl, KO and ALKKO mice as well as ALK tumors were analyzed by Western blot analysis. Tubulin served as loading control. Analysis was performed in biological triplicates. Asterisks indicate unspecific band.

### *Dnmt1* deletion reduces proliferation of ALKKO cells

We next studied the proliferative capacity of ALKKO thymocytes compared to the other genotypes by assessing Ki67 expression. As expected, ALK tumor cells showed significantly higher proliferation rates as compared to thymocytes of Ctrl, KO and ALKKO mice (Fig. 4A). A trend towards lower proliferation compared to Ctrl and KO was detectable in ALKKO thymi. In order to specifically analyze ALK expressing cells, we next performed co-staining of ALK and Ki67 in ALK and ALKKO tissues using immunofluorescence (Fig. 4B). Quantification of ALK and Ki67 double-positive cells revealed that ALK expressing cells showed a significant reduction of Ki67 positivity in ALKKO thymi compared to ALK tumors, with 85,7% double positive in ALK versus 58,7% in ALKKO samples. In particular, ALKKO cells with high ALK expression levels showed low expression of Ki67, whereas cells with high Ki67 levels showed very low expression of ALK, indicating that ALK cells cannot induce or maintain stable proliferation upon *Dnmt1* deletion.

**Figure 4.**
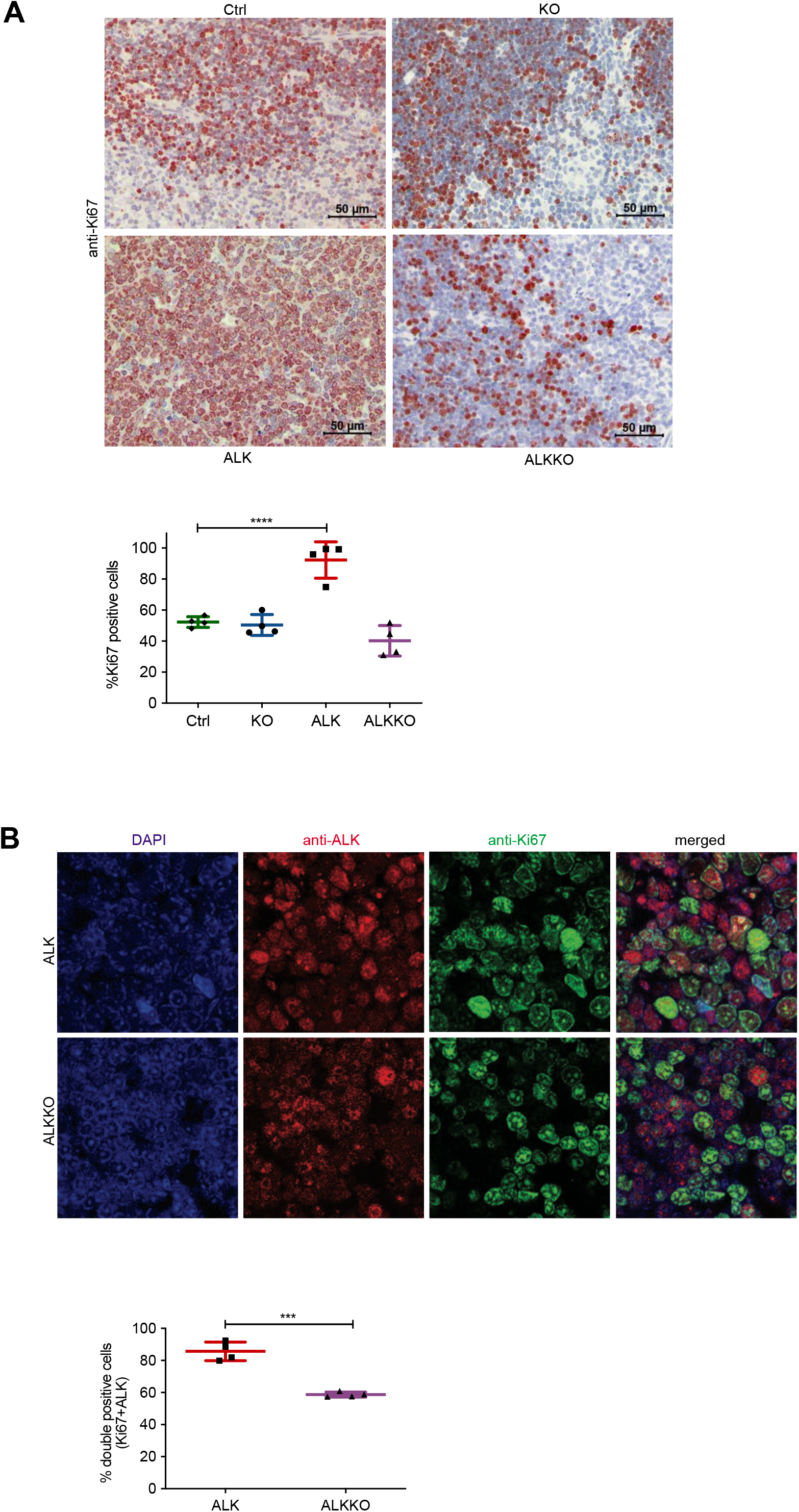
Deletion of *Dnmt1* results in reduced proliferation of ALK+ cells. **A,** Cell proliferation analysis by immunohistochemistry staining of Ctrl, KO, ALKKO thymi and ALK tumor tissues using Ki67 antibody. The graph shows quantification of Ki67 positive cells using Definiens TissueStudio 4.2 software. Non-proliferative areas in the thymus were excluded from analysis. Data are shown as mean±SD, ***p<0.001, pair-wise comparison to control using unpaired t-test, n=4. **B,** Double immunofluorescence staining of ALK tumors and ALKKO thymi. Tissues were stained with antibodies against ALK (red) and Ki67 (green) and counterstained with DAPI (blue). Pictures were acquired with identical pixel density, image resolution and exposure time. The graph shows quantification of immunofluorescence staining by counting Ki67/ALK double-positive relative to total number of cells (DAPI positive) of two equally sized areas per tumor/thymus from four biological replicates, respectively. Cell counting was done by two individuals and slides were blinded for counting. Data are shown as mean±SD, ***p<0.001, using unpaired t-test.

### *Dnmt1* knockout inhibits ALK-dependent transcription programs

To investigate the molecular mechanisms associated with *Dnmt1* deletion in ALK transgenic mice, we performed RNA-seq analyses of thymocytes isolated from Ctrl, KO and ALKKO mice and compared it to our tumor RNA-Seq data (Fig. 5). Sample distance and principal component analysis revealed a clear separation of ALK tumors from all other genotypes, which were clustering together (Fig. 5A and Supplementary Fig. 5A). Within the groups, ALK tumors showed the largest heterogeneity. Along these lines, unsupervised clustering of the 5% most variably expressed genes revealed a clear separation of tumor samples from all other genotypes, which showed highly similar expression patterns (Fig. 5B). Differential gene expression analysis showed that ALKKO displayed highly similar gene expression patterns compared to Ctrl cells: We only found 8 genes to be significantly down- and 99 genes upregulated (Fig. 5C, Supplementary Fig. 5B and Supplementary Table 2). When compared to ALK tumor samples, ALKKO was similar to KO and Ctrl samples and showed a high degree of deregulation with 2819 genes up- and 1355 genes downregulated (Supplementary Fig. 5B).

**Figure 5.**
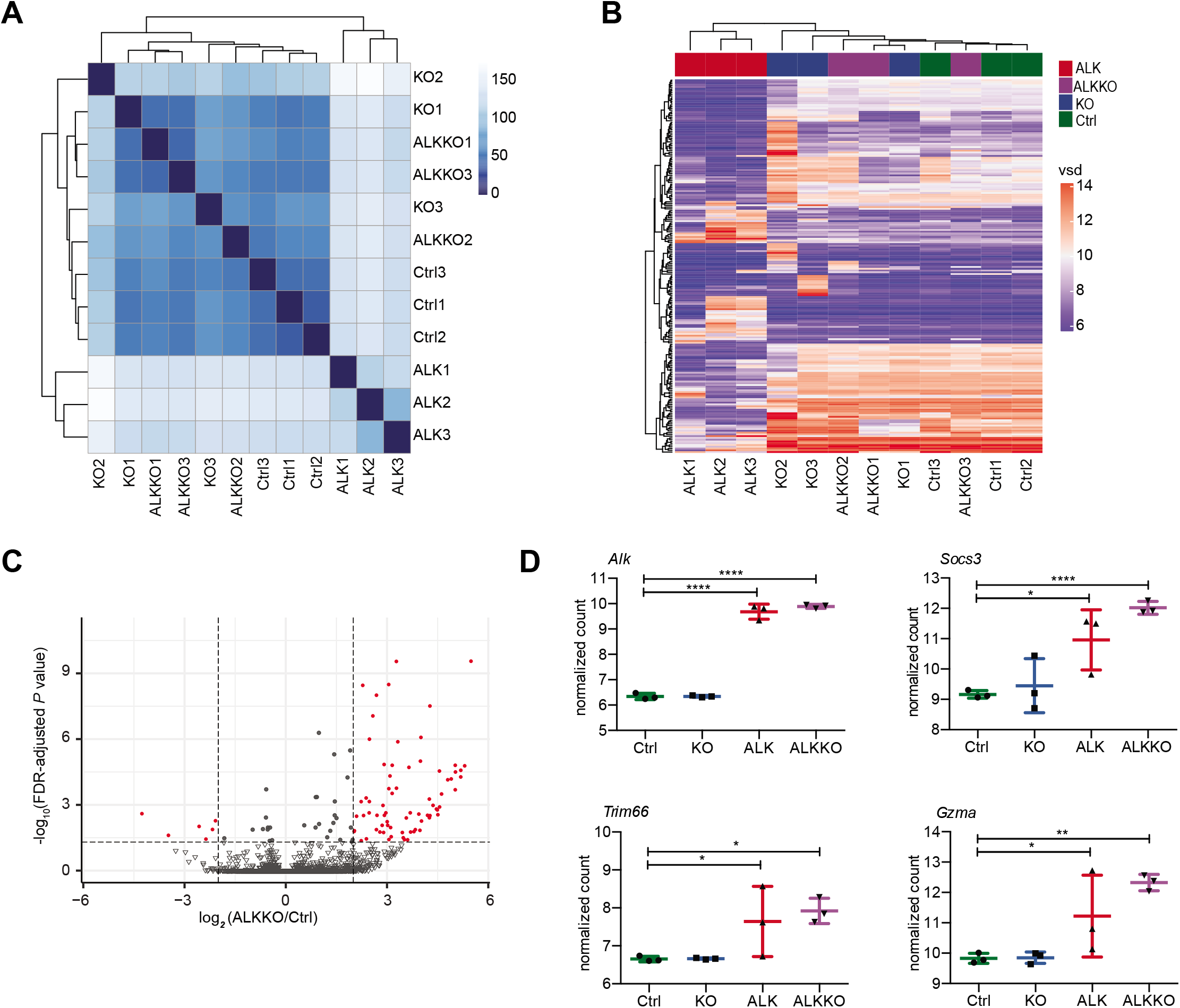
High similarity in gene expression between ALKKO and Ctrl thymocytes. **A,** Hierarchical clustering heatmap illustrating sample to sample Euclidian distances based on variance stabilizing transformations (VST) of RNA-seq gene expression values of all genes of individual Ctrl, KO, ALKKO thymi and ALK tumor samples. **B,** Heatmap showing unsupervised clustering of the top 5% most variable genes among all samples in Ctrl, KO, ALK and ALKKO using variance stabilizing data (vsd). **C,** Volcano plot displaying the significant differences in gene expression between ALKKO compared to Ctrl thymocytes, where red dots indicate significant differentially expressed genes (FDR<0.05, absolute log_2_(FC)>1) and grey scale dots show non-significant differentially expressed genes, between this two groups. **D,** Gene expression levels of *Socs3, Trim66* and *Gzma* based on normalized counts from RNA-seq analysis of ALK tumor and ALKKO thymus samples compared to Ctrl and KO thymi. Data are represented as mean±SD, *p<0.05, **p<0.01, ****p<0.0001, using ordinary one-way ANOVA followed by multiple comparison using Fisher’s least significant difference (LSD) test.

We were particularly interested in a small group of genes, which were consistently deregulated in both ALK+ cell types relative to Ctrl and *Dnmt1* KO cells, as identified by pairwise comparisons of gene expression differences, since they might constitute direct targets of NPM-ALK upstream of DNMT1 (Supplementary Fig. 5C). Apart from ALK, we found 7 genes *(Tha1, Trim66, Gzma, Socs3, Gm5611, 5830468F06Rik, Gm17910),* which were upregulated both in ALK and ALKKO cells compared to Ctrl.

These include the suppressor of cytokine signaling 3 *(Socs3),* which is a regulator of the JAK/STAT signaling pathway and was also found upregulated in human ALK+ ALCL cell lines (53), tripartite motif containing 66 *(Trim66),* which is part of the RAS pathway that regulates DNMT1 expression and is known to promote proliferation (54–56) and granzyme A *(Gzma)* a canonical cytotoxic gene that is involved in cancer initiation and progression (57) (Fig. 5D).

These data suggest that expression of ALK initially affects a small number of regulatory genes including *Socs3*, *Trim66* and *Gzma*, while a downstream substantial rewiring of the whole transcriptional program eventually accompanies induced tumorigenesis.

### Changes in DNA methylation following *Dnmt1* deletion

In order to assess the effect of DNMT1 depletion on global DNA methylation in the different genotypes, we analyzed global and site-specific CpG methylation, using dot blot analysis measuring 5-methyl Cytosine (5mC) levels and genome-wide methylation patterns using reduced representation bisulfite sequencing (RRBS), respectively (Fig. 6A,B and Supplementary Fig. 6A). As shown previously (52), KO samples revealed a slight but not significant decrease in global DNA methylation compared to Ctrl samples. In the double mutant, ALKKO, we observed a significant decrease in global DNA methylation levels as compared to Ctrl samples. The loss of DNA methylation in KO and ALKKO samples was accompanied by strong induction of *IAP* retrotransposon transcription, which was especially high in ALKKO, in line with higher loss of global methylation in those samples (Supplementary Fig. 6B).

**Figure 6.**
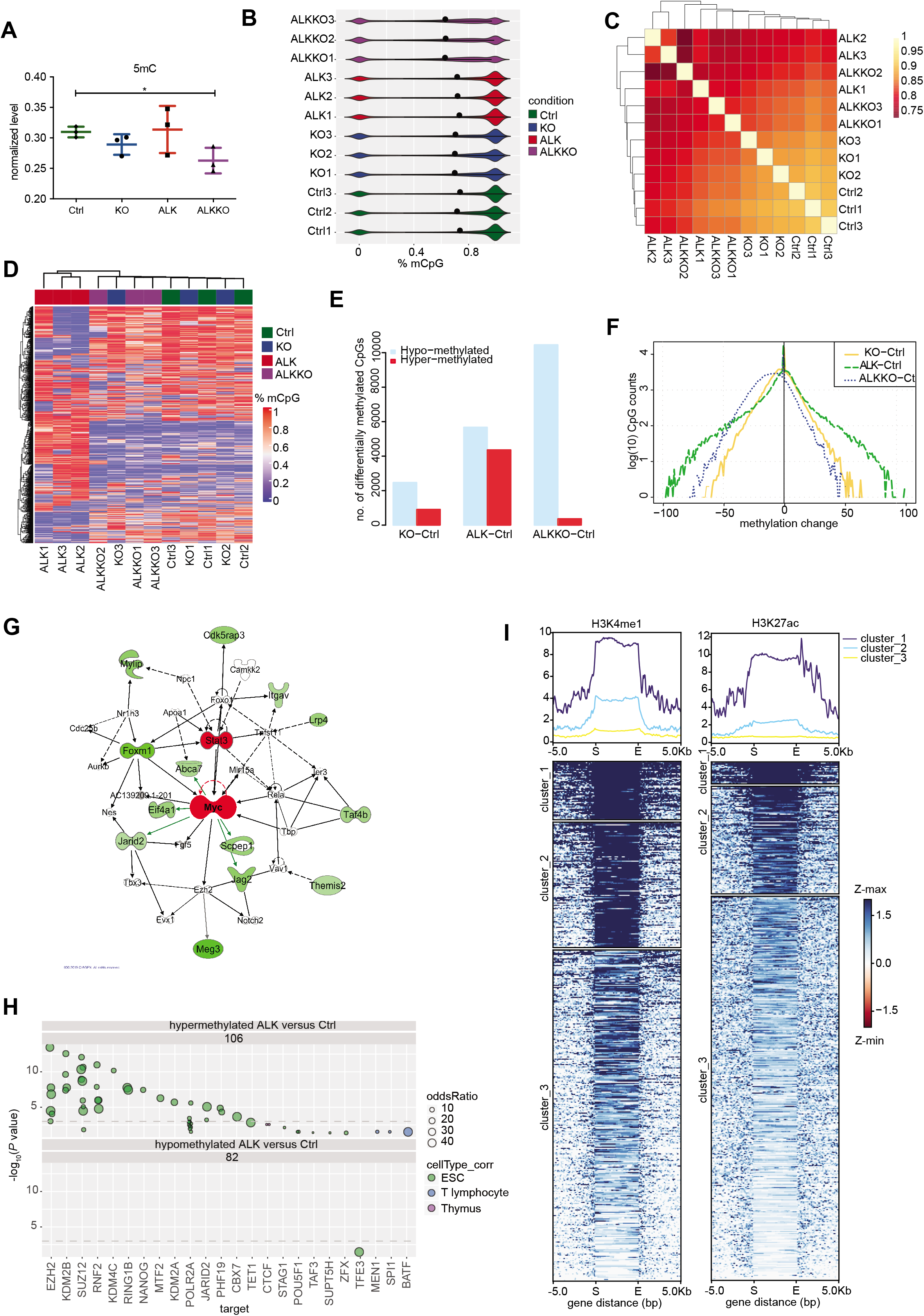
DNA methylation changes in ALK tumors and *Dnmt1* knockout thymi. **A,** Quantification of global DNA methylation levels by dot blot analysis using 5mC immunodetection. 5mC signal intensities were normalized to total DNA input based on methylene blue staining. Data are represented as mean±SD, * p<0.05, using one-way ANOVA, followed by unpaired t-test, n=3. **B,** Violin plots indicating the bimodal distribution of methylation levels determined by reduced representation bisulfite sequencing (RRBS) in three biological replicates of Ctrl, KO and ALKKO thymi as well as ALK tumors. Shown are percent methylation per CpG (%mCpG). Black dots indicate the median percentage of mCpG in each sample. **C,** Correlation heatmap between individual Ctrl, KO, ALKKO and ALK samples based on DNA methylation levels of single CpGs showing correlation coefficients between samples. **D,** DNA methylation heatmap using unsupervised clustering of the top 1% most variable CpGs in Ctrl, KO, ALK and ALKKO samples. The color code represents mean methylation levels of single CpGs. **E,** Significantly hyper- (red) and hypo- (blue) methylated CpGs resulting from comparison of KO, ALKKO and ALK versus Ctrl samples identified by methylKit analysis (p<0.01; b-value difference >25%). **F,** Methylation changes of individual CpGs relative to Ctrl for all genotypes (as in E) shown as density plot. Numbers on the y-axis are log(10) CpG counts. **G,** Network analysis based on Ingenuity^®^ Pathway Analysis of significantly hypomethylated promoter regions in ALK versus Ctrl samples. Significantly hypomethylated genes are depicted in green, upstream regulators are depicted in red. **H,** Locus overlap analysis (LOLA) region set enrichment analysis for differentially methylated CpGs between transcriptional subtypes (binned into 1-kilobase tiling regions). The plot shows region sets from embryonic stem cells (ESC, green), T lymphocytes (blue) and thymus (purple) with p<0.05. **I,** Differentially methylated regions (DMRs) between ALK versus Ctrl samples were used to map histone modifications (H3K4me1 and H3K27ac) of ENCODE regions defined by ChIP-seq data of thymus samples using *#*-means clustering (*ł*=3). The heatmap indicate overlap of individual DMRs expanded to 5kb on both sites with H3K4me1 (left) and H3K27ac (right) in three distinct clusters, based on different signal intensity. Gene distance indicates predicted enhancers mapped within at a 5-kb window indicating start (S) and end (E) of the DMR region. Z-min/max show the intensity of the H3K4me1 and H3K27ac ChIP-seq signals.

CpG methylation showed the highest correlation between Ctrl and KO samples, whereas both ALK and ALKKO samples showed lower correlations in relation to other genotypes but also amongst the replicates (Fig. 6C). Similarly, unsupervised hierarchical clustering of the top 1% most variable CpGs indicated a separation of ALK tumor samples compared to Ctrl, KO and ALKKO thymi and a closer proximity between Ctrl and KO compared to ALKKO samples, suggesting that the largest changes in methylation occur in the tumor samples compared to the other genotypes (Fig. 6D).

Compared to Ctrl, ALK samples showed both hyper- and hypomethylation, whereas the KO and ALKKO samples were enriched for hypomethylated sites (Fig. 6E). When comparing the change in methylation per site, we observed that in ALKKO compared to Ctrl most sites lost less than half of their methylation, whereas in ALK versus Ctrl we detected CpGs with particularly large changes reflecting stable tumor-specific changes in DNA methylation patterns (Fig. 6F).

Annotation of differentially methylated CpGs revealed that ALK tumor samples mainly gained DNA methylation in promoter regions, CGIs and shores, whereas intergenic and inter-CGI sites preferentially lost methylation (Supplementary Fig. 6C,D). The large number of hypomethylated CpGs between Ctrl and ALKKO was mainly associated with intergenic and inter-CpG island regions and to a lesser extent with upstream promoter regions, exons, shores and shelves (Supplementary Fig. 6E,F).

In order to identify potential tumor relevant pathways associated with differentially methylated regions, we performed ingenuity pathways analysis (IPA^®^, www.qiagen.com/ingenuity), interrogating promoters that showed tumor-specific hypomethylation. Interestingly, those analyses identified c-MYC as a significant upstream regulator and several genes connected to the cellular c-MYC network were hypomethylated in ALK tumor samples (Fig. 6G). Among these genes, *Cdk5rap3, Lrp4, Eif4a1* and *Taf4b* were also significantly upregulated in tumors compared to Ctrls based on RNA-seq data. Interestingly, all four genes have been implicated in proliferation control and tumorigenesis before (58–61), highlighting their potential relevance for ALK driven lymphomagenesis.

To get further information about the genomic regions that are affected by aberrant methylation in ALK tumor cells we performed genomic region enrichment analysis on differentially methylated CpGs tiled into 1kb genomic regions using locus overlap analysis (LOLA) (62). We used published datasets integrated in the LOLA software tool including murine embryonic stem cells (ESC), T lymphocytes and thymus for comparison with our DNA methylation profile of ALK tumor cells. Hypermethylated CpG regions in ALK tumor cells showed strong enrichment of binding sites for proteins associated with chromatin remodeling in ESC, in particular EZH2, SUZ12, MTF2, JARID2 and PHF19, which are associated with polycomb repressive complex 2 (PRC2). In addition, we detected ESC specific binding of the histone demethylase KDM2B, members of polycomb repressive complex 1 (PRC1; RNF2, CBX7) as well as the DNA demethylase TET1 to be enriched in hypermethylated regions in ALK tumors (Fig. 6H). Epigenetic switching from polycomb repressive complex (PRC) marks to DNA methylation is a well-described phenomenon in human tumor cells reducing epigenetic plasticity of tumor cells (9), (63). Furthermore, T lymphocyte specific binding of the tumor suppressor MEN1, which has a role for Th2 cell function and prevents CD8+ T cell dysfunction (64,65), SPI1, a master regulator of early T cell transcription (66) and BATF, a AP-1 family member, which has been implicated in human ALCL growth and survival (67) were significantly overlapping with hypermethylated sites in ALK mouse tumors. For hypomethylated regions, we detected an overlap of ESC specific TFE3 binding sites. TFE3 has been implicated in Wnt signaling (68) and thymus-dependent humoral immunity (69).

Enhancers play an important role in regulating gene expression during development and it has been shown that enhancer methylation can be drastically altered in cancer, which can be associated with altered expression profiles of cancer genes (70). Therefore, we compared differentially methylated regions (DMRs), defined over 1kb tiling windows, between ALK tumor cells and Ctrl thymocytes to ENCODE ChIP-seq datasets for active enhancers including monomethylation of histone H3 lysine K4 (H3K4me1) and acetylation of histone H3 lysine 27 (H3K27ac) in murine thymi (71,72). Among 470 DMRs tested, we detected a specific overlap of 118 DMRs with H3K27ac and 167 DMRs with H3K4me1, respectively (Fig. 6I). Of these, 95 DMRs overlapped with both H3K4me1 and H3K27ac, indicative of active enhancers in thymocytes (Supplementary Table 3). Interestingly, nearby genes of several differentially methylated enhancers showed significantly deregulated gene expression (Supplementary Table 3). Among those genes we found *Runx1*, a critical transcription factor for early T cell development of thymic precursors and T cell maturation (73,74), or *Socs3*, which was also implicated in human ALCL (75,76). Downregulated genes included the phosphatase and tumor suppressor *Ptpn13* (77) and the TGF-beta ligand *Bmp3,* which might result in altered TGF-beta signaling as observed in human ALK-positive cancers (78).

Together these data suggest that NPM-ALK driven transformation is accompanied by CGI hypermethylation and hypomethylation of intergenic regions, which is reminiscent of epigenetic reprogramming events in human tumors and affects major oncogenic signaling pathways. Furthermore, analysis of regions that are associated with differentially methylated CpGs in ALK tumor cells compared to Ctrl thymocytes revealed hypermethylation of binding sites associated with PRC1 and PRC2 components and T cell specific transcription factors and a significant overlap with thymus specific enhancers.

### Loss of collaborative methylation patterns in tumors

Recent literature has challenged the standard model of DNA methylation inheritance, in which CpGs are presumed as independent, but has rather suggested a collaborative model that takes methylation levels of neighboring CpGs into account (79,80). According to this model, which also considers the distances of neighboring sites, methylated CpGs would enforce methylation of adjoining CpGs, whereas nonmethylated CpGs would induce rather unmethylated states depending on the presence of DNMTs and TET enzymes. We therefore analyzed whether the distance between two neighboring CpGs had an influence on methylation changes of those CpGs in ALK tumor or knockout cells (ALK and ALKKO) relative to Ctrls (Fig. 7A). ALK tumor cells showed stronger hypermethylation in nearby CpGs (0-5bp distance), most likely reflecting tumor specific CGI hypermethylation. More distant sites showed equal gains and losses of methylation in the ALK samples. Further, we found that in the KO samples as well as in the double mutant ALKKO, highest methylation loss was observed at CpGs more distant from neighboring CpGs, suggesting that DNMT1 is needed for methylation maintenance of distant CpGs.

**Figure 7.**
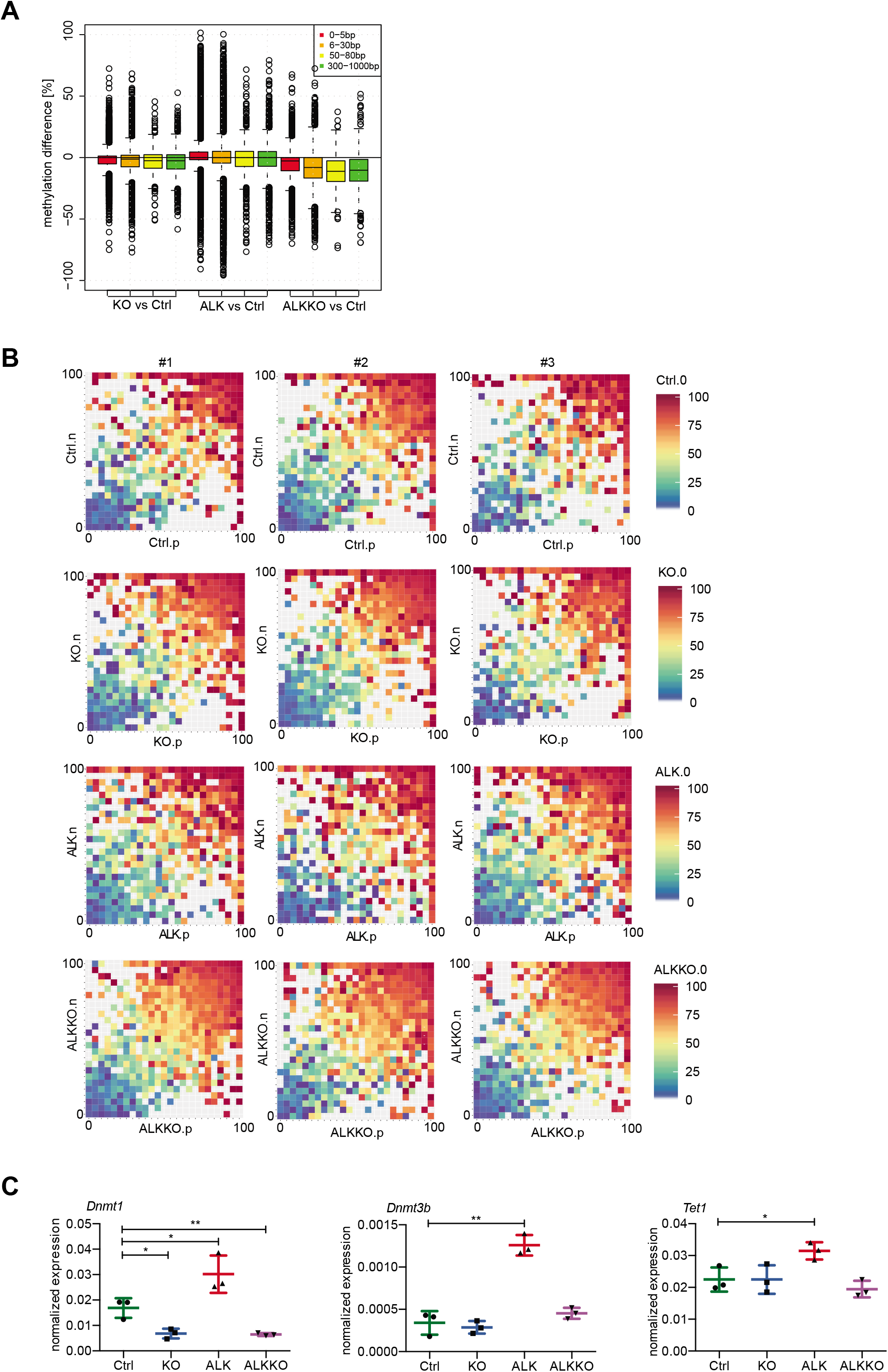
Analysis of collaborative DNA methylation. **A,** Methylation difference of neighboring CpGs relative to Ctrls in KO, ALK and ALKKO samples as analyzed based on the distance between CpGs. **B**, CpG triplet analysis of neighboring CpGs within a maximal distance of 20 bp. Each square represents three neighboring CpGs. The color of the squares indicates the methylation level of the middle CpG from unmethylated (blue) to fully methylated (red). The x and y axes represent the methylation levels of the two neighboring CpGs annotated as (n) and (p). Squares close to the diagonal indicate highly correlated CpG triplets, whereas dispersed squares represent non-correlated triplets. **C,** RT-qPCR of *Dnmt1, Dnmt3b* and *Tet1* expression Ctrl, KO and ALKKO thymocytes as well as ALK tumor cells normalized to *Gapdh* expression. Analyses were performed in biological triplicates. Data are represented as mean±SD, *p<0.05, **p<0.01, pairwise comparison to the Ctrl unpaired t-test.

We further examined closely associated CpG triplets, defined as CpGs with neighboring CpGs within less than 20bp distance to both sites, in three biological replicates of each genotype (Fig. 7B). We found that in Ctrl samples methylation levels between neighboring sites are highly correlated, in a way that a highly methylated CpG is flanked by highly methylated CpGs, whereas unmethylated CpGs are flanked by unmethylated CpGs. Thus, in a correlation plot, most CpGs are found close to the diagonal in Ctrl samples (Fig. 7B). *Dnmt1* knockout samples showed a general trend towards lower methylation but retained the correlation among the CpG triplets, as illustrated by the vicinity of CpG triplets to the diagonal in KO and ALKKO samples, suggesting that DNMT1 is not required to warrant cooperative methylation. Interestingly, cooperative DNA methylation seems to be disturbed in ALK tumor cells, where we detected a loss of correlation and observed a higher methylation difference of neighboring CpGs, indicating a loss of collaboration between those sites.

Together, these data suggest that heterogeneous DNA methylation patterns in ALK+ tumors are characterized by low correlation of DNA methylation between nearby CpGs, which might be associated with the deregulation of methylation controlling enzymes including *Dnmt1*, *Dnmt3b* and *Tet1* (Fig. 7C) that we observed based on RT-qPCR analyses. Interestingly, the correlation between neighboring CpGs is maintained upon DNMT1 depletion, suggesting that *de novo* methyltransferases such as DNMT3a/b can compensate and control collaborative DNA methylation at sites in close vicinity.

## DISCUSSION

The ALK protein has been identified as a driver oncogene in diverse cancers, based on its overexpression following genetic translocation, amplification or mutation (81–83). Overexpression of the human NPM-ALK oncogene in mouse T cells results in lymphomagenesis with 100% penetrance. In this model, the relatively long period of latency despite NPM-ALK expression and activation of downstream targets such as STAT3 and DNMT1, suggests that a secondary event is necessary for transformation and tumorigenesis. Accordingly, the NPM-ALK translocation and its transcripts can be detected in peripheral blood cells of human healthy donors implying that a single genetic event is not sufficient to induce transformation (84). Both our transcriptomic and epigenomic data point towards MYC as the potential secondary driver in this model. MYC was previously implicated in different ALK dependent malignancies including non-small cell lung cancer (NSCLC) neuroblastoma and ALK+ ALCL (35,85–90). Furthermore, MYC was shown to directly regulate expression of *Cdk4* and *Cdk6*, affecting cell cycle progression at multiple points (91). We observed a gradual upregulation of those genes on mRNA as well as protein level from tumor initiation to end-stage tumors. Correspondingly, we observed strong deregulation of *Notch1* in the transgenic mouse model. The NOTCH-MYC axis plays a major role in the development of T cell acute lymphoblastic leukemia (T-ALL), resulting from the transformation of immature T cell progenitors (92). This suggests that similar processes are driving ALK dependent transformation, resulting in hyperproliferation and transformation of thymocytes. Notably, deletion of *Dnmt1* in a MYC-driven T cell lymphoma model delayed lymphomagenesis and resulted in reduced proliferation of tumor cells (93).

During lymphomagenesis, we observed a gradual decrease of DP thymocytes subsets and a corresponding increase of DN and immature CD8 SP (i.e. TCRβ-) cells. Similarly, human ALK+ ALCL display a progenitor cell signature as indicated by epigenomic profiling and a subgroup of ALCL might arise from innate lymphocyte cells as recently described based on transcriptomic analyses (46,67). The fact that we observe NPM-ALK expressing tumor cells that show a surface marker expression pattern that is typical for DN cells, as well as the presence of ISP cells suggests, that an immature thymic cell population is targeted for transformation or that a more mature cell population regresses to a progenitor stage during transformation. Again, MYC might be central to this event, in line with data showing that overexpression of MYC, activated AKT and inhibition of intrinsic apoptosis by expression of BCLXL results in rapid transformation of mature CD4 and CD8 SP mouse T cells (94). Expression of reprogrammed DN lymphoma stem cells was recently described for a Lck-dependent NPM-ALK mouse model (95).

The elevated expression of DNMT1 appeared to be an early event following NPM-ALK induction in our model and depletion of DNMT1 completely abrogated lymphomagenesis. Intriguingly, we found DNA methylation signatures, highly resembling human tumors (96), with characteristic CGI hypermethylation and genomewide hypomethylation as well as DNA hypermethylation of polycomb repressive marks (9). Generally, epigenomic patterns appeared to be highly heterogeneous between individual ALK tumors and showed a loss of cooperative CpG methylation. These observations went hand in hand with upregulation of DNMTs (DNMT1, DNMT3b) and TET1, implying that the oncogenic driver NPM-ALK has the potential to interfere with methylation homeostasis and to induce stochastic methylation aberrations. Interestingly, deletion of DNMT1 did not interfere with cooperative DNA methylation between closely neighboring CpGs, since KO and ALKKO cells maintained high correlation between triplet CpGs with a CpG distance less than 20 bp. Thus, we propose that aberrant DNA methylation patterns in tumors might not fully depend on DNMT1 deregulation, and that methylation cooperativity at close-by CpGs is independent of the maintenance machinery, but rather dependent on DNMT3a/b and TET enzymes. In general, DNMT3a and DNMT3b primarily bind to methylated CpG-rich regions, however DNMT3b seems to exhibit additional preferences for actively transcribed genes and correlates with H3K36me3 marks (97). In addition, DNMT3a/b are important to counteract global DNA methylation loss due to imperfect DNMT1 fidelity during DNA replication in mice (98). *De novo* methylation occurs more frequently at adjacent CpGs in a distance dependent manner (97), further indicating that collaboration between neighboring CpGs could be maintained through DNMT3a and DNMT3b in KO and ALKKO samples.

In summary, we conclude that ALK dependent oncogenic pathways result in deregulation of genome-wide DNA methylation patterns during tumorigenesis, that affect important regulatory regions including lineage specific transcription factor binding sites and enhancers of T cell specific genes and tumor suppressors. Additionally, we suggest that the deregulation of key epigenetic enzymes is a prerequisite to enable tumor formation and loss of maintenance of tumor specific DNA methylation patterns results in a proliferation block and lack of T cell transformation.

## METHODS

### Mice

Transgenic mice carrying the human NPM-ALK fusion-gene under the T cell specific *Cd4* enhancer-promoter system were crossed with mice carrying a conditional T cell specific deletion of *Dnmt1 (Cd4-Cre* driven recombinase) (31,52,99). The NPM-ALK mice were obtained from Lukas Kenner, Department of Pathology, Medical University of Vienna, and the *Dnmt1* knockout mice were obtained from Christian Seiser, Center for Anatomy and Cell Biology, Medical University of Vienna. The genetic background of mice was mixed (C57Bl/6xSV/129). Mice were kept under specific pathogen-free (SPF) conditions at the Center for Biomedical Research, Medical University of Vienna and experiments were done in agreement with the ethical guidelines of the Medical University of Vienna and after approval by the Austrian Federal Ministry for Science and Research (BMWF; GZ.: 66.009/0304.WF/V/3b/2014).

For genotyping, tissue samples obtained from ear clipping were incubated with tail lysis buffer (100 mM Tris pH 8.0, 5 mM EDTA pH 8.0, 200 mM NaCl, 0.1% SDS) and 40 μl of proteinase K (10 mg/ml) o/n at 56°C. The next day, 170 μl of 5M NaCl were added, the solution was centrifuged for 10 min at maximum speed and the supernatant was transferred to a new tube. 500 μl of isopropanol were added and after centrifugation and washing with 70% EtOH the DNA was dried and dissolved in 150 μl of sterile water. Genotyping was performed with Promega GoTaq Mastermix according to manufacturer’s suggestions.

### FACS analysis

Single cell suspensions of thymus, tumor and spleen were obtained by passaging the tissues through a 70-μm nylon cell strainer in staining buffer (PBS supplemented with 2% FCS and 0.1% sodium azide). 3 x 106 cells were incubated for 5 min on ice with Fc-block (Pharmingen), following incubation with cell surface markers for 30 min on ice. Cells were washed and fixed with eBioscience^TM^ FoxP3 Transcription Factor Fixation/Permeabilization Solution for 1 hour at 4°C before. Intracellular staining for ALK (D5F3, CST #3633) was done for 1 hour at 4°C followed by incubation with Alexa Fluor^®^ 647 goat anti-mouse IgG antibody (Thermo Fisher, Catalog # A-21244) for another 1 hour at 4°C. Cells were washed and resuspend in 100 μl staining buffer. Cells were measured with a BD Fortessa flow cytometer and analyzed using FlowJo software.

### RNA and DNA isolation from murine tumor and thymic tissues

RNA and DNA were isolated using the QIAGEN AllPrep DNA/RNA mini isolation kit to enable simultaneous isolation of nucleic acids from the same specimen. Tissues were homogenized using EPPI-Mikropistills (Schuett-biotec) and RNA and DNA isolation was performed according to the manufacturer’s protocol. RNA and DNA were eluted in nuclease free ddH_2_O.

### Protein extraction and Western Blot

For protein extraction from tumors and thymi, the tissue was dounced and homogenized in lysis buffer as previously described (100). Protein concentrations were measured using Bradford and 20 μg were used for analysis by SDS-page and Western blot as previously described (100). The following antibodies were used for protein expression analysis: ALK (D5F3, CST #3633), pALK (Tyr1278, CST #6941), STAT3 (D3Z2G, CST #12640), pSTAT3 (Tyr705, CST #9145), DNMT1 (H300, sc-20701), CDK4 (Santa Cruz, C-22, sc-260), CDK6 (Sigma-Aldrich, HPA002637), c-MYC (fE5Q6W, CST #18583), PCNA (Abcam, ab2426) and alpha-TUBULIN (1E4C11, Proteintech, 66031-1-Ig). Goat anti-rabbit IgG HRP conjugate, JD111036047 and rabbit anti-mouse IgG HRP conjugated, JD315035008 antibodies were used as secondary antibodies.

### Quantitative reverse transcription PCR (RT-qPCR)

For RT-qPCR, RNA was isolated as described above and 1 μg of RNA was used for random hexamer cDNA synthesis using the qScript cDNA Synthesis Kit (Quantabio) according to supplier’s protocol. cDNA was diluted to a final concentration of 5 ng/μl and RT–qPCR was performed with KAPA SYBR^®^ FAST qPCR kits (KAPA Biosystems) on a C1000 thermal cycler, CFX96 real-time system (Biorad) using 10 ng of cDNA per reaction. Three biological replicates per genotype were processed. GAPDH was used for normalization. The following primers were used for RNA-sequencing validation:

*Gapdh* fw: 5’-CGACTTCAACAGCAACTCCCACTCTTCC-3’
*Gapdh* rv: 5’-TGGGTGGTCCAGGGTTTCTTACTCCTT-3’
*Cd44* fw: 5’-ATGAAGTTGGCCCTGAGCAA-3’
*Cd44* rv: 5’-GTGTTGGACGTGACGAGGAT-3’
*Dnmt1* fw: 5’-AGGAGAAGCAAGTCGGACAG-3’
*Dnmt1* rv: 5’-CTTGGGTTTCCGTTTAGTGG-3’
*Notch1* fw: 5’-TGGCAGCCTCAATATTCCTT-3’
*Notch1* rv: 5’-CACAAAGAACAGGAGCACGA-3’
*Myc* fw: 5’-AGTGCTGCATGAGGAGACAC-3’
*Myc* rv: 5’-GGTTTGCCTCTTCTCCACAG-3’
*Cdk4* fv: 5’-TCCCAATGTTGTACGGCTGA-3’
*Cdk4* rv: 5’-ACGCATTAGATCCTTAATGGTCTCA-3’
*Cdk6* fw: 5’-CAGCAACCTCTCCTTCGTGA-3’
*Cdk6* rv: 5’-GATCCCTCCTCTTCCCCCTC-3’
*IAP* fw: 5’-ACTAAcTCCTGCTGACTGG-3’
*IAP* rv: 5’-TGTGGCTTGCTCATAGATTAG-3’

### Immunohistochemistry

Tumor and thymic tissues were fixed in 4% paraformaldehyde, dehydrated in ethanol and embedded in paraffin. 2 μM sections were cut, attached to slides, dewaxed and rehydrated. Epitopes were retrieved by heat-treatment in citrate buffer (pH 6.0, DAKO) or Tris-EDTA buffer (pH 9.0, DAKO). Slides were processed and counterstained as previously described (100). Primary antibodies against DNMT1, ALK pSTAT3 (listed above) or Ki67 (eBioscience, 14-5698-80) were used for staining.

Pictures were taken with a Zeiss Axio10 (Zeiss) microscope and a Gryphax camera (Jenaoptics) and quantification of positive cells was performed using Definiens Tissue Studio 4.2 Software (Definiens Inc.). Stainings were performed in 4 biological replicates for each genotype. For each biological replicate 4 representative pictures were analyzed and counts were averaged. For Ki67 stainings slides were scanned using the Pannoramic 250 Flash III scanner (3DHISTECH) and analyzed using the Definiens software. Non-proliferative areas in thymi were excluded from quantification.

Quantification results are shown as means ± SEM. The significance of the differences between mean values was determined by one-way ANOVA followed by pair wise comparisons to the control group using unpaired *t*-tests.

### Immunofluorescence

Tumor and thymic tissues were formalin fixed and paraffin embedded as described above and epitopes were retrieved by heat-treatment in citrate buffer (pH 6.0, DAKO). Slides were washed in 0.1% PBS-Tween20 (PBS-T), permeabilized in 0.3% TritonX-100 in PBS-T and blocked in blocking solution (10% goat serum in 0.1% TritonX-100 in PBS-T). Primary antibodies against ALK (CST #3363) and Ki67 (eBioscience, 14-5698-80) were diluted 1:250 and 1:1000 in blocking solution and incubated at 4°C overnight. After incubation with secondary antibodies (Alexa-Fluor 594 goat antirabbit, Invitrogen Cat # A-11012 and Alexa-Fluor 488 goat anti-rat, Invitrogen Cat # A-11006) diluted 1:1000 in blocking solution, slides were counterstained with DAPI (1:50000 from 10 mg/ml stock solution, Serva Eletrophoresis) and embedded with geltol (Calbiochem). 2 representative pictures of 4 biological replicates per genotype were taken with LSM 5 Exciter (Zeiss) with same exposure time for all slides and quantified by blinding pictures and counting of single positive and double positive cells in 2 3×3 cm squares per picture. Quantification results are shown as means ± SEM. The significance of the differences between mean values was determined by one-way ANOVA followed by pair wise comparisons to the ALK group using unpaired *t*-tests.

### Dot blot analysis of methylated DNA

Genomic DNA was isolated as described above and diluted to a final concentration of 250 ng. DNA was denatured at 100°C for 10 min in 0.4M NaOH and 10 mM EDTA solution and neutralized using 2M ice cold ammonium acetate, pH 7.0 before it was applied to the dot blot apparatus to spot the DNA on a nitrocellulose membrane, presoaked in 6x SSC buffer. After washing the membrane in 2x SSC buffer the DNA was UV-crosslinked using (UV Stratalinker^®^ 2400, Stratagene). Subsequently the membrane was blocked for 1 hour at room temperature with 5% milk in PBS-T and incubated with the primary antibody directed against 5mC (D3S2Z, CST #28692) overnight at 4°C. After incubation with the secondary antibody (goat anti-rabbit IgG HRP conjugate, JD111036047) 5mC signal was detected using the ChemiDoc^TM^ XRS+ Imaging System (Biorad) and analyzed using Image Lab^TM^ Software (Biorad). Methylene blue staining (0.02% methylene blue in 0.3M sodium acetate) was used for an internal DNA loading control. Membranes were incubated in methylene blue staining solution for 10 min and membranes were destained 3x 10 min in water. Methylene blue staining was measured and analyzed as described above. Level of 5mC was calculated as a ratio of 5mC to methylene blue signal intensity in 3 biological replicates per genotype and 3 technical replicates, respectively.

### RNA-Sequencing (RNA-Seq)

RNA and DNA were isolated as described above. RNA concentrations were measured on the Nanodrop 2000 (Invitrogen) and 1000 ng were sent to the Biomedical Sequencing Facility (BSF, CeMM, Vienna). RNA integrity was tested using the Agilent Bioanalyzer. Stranded mRNA-seq (poly-A enrichment) library preparation was performed and sequenced employing an Illumina HiSeq3000/4000 platform (50 nucleotide single-end reads).

### RNA-seq data analysis

Reads were quality-controlled using fastQC (101) and pre-processed using trimgalore (http://www.bioinformatics.babraham.ac.uk/projects/trim_galore/) to trim adapter and low quality sequences. The reads were aligned to mouse genome (mm10) and processed further using STAR (102). Differential gene expression levels of the transcripts were quantified by HTSeq (103) and analyzed using the Bioconductor package DESeq2 (104). Genes with an FDR-adjusted p-value below 0.05 and a absolute fold change of 2 were considered significantly differentially expressed.

To gain insight into the nature of differentially expressed genes in each analysis, the gene set enrichment analysis (GSEA) of significantly deregulated genes between ALK tumors and Ctrl thymi was performed using oncogenic signature gene sets from MSigDB (33).

### Reduced representation bisulfite sequencing (RRBS)

DNA was isolated as described above. DNA concentration was measured using Qubit^TM^ dsDNA HS Assay Kit from Invitrogen according to manufacturer’s protocol. In total 500 ng (25 ng/μl, 20 μl total) of DNA were sent for RRBS analysis (BSF, CeMM, Vienna) according to established protocols (105,106).

### RRBS data analysis

The global changes in methylation, individual CpGs and clusters of covered CpGs were analyzed using packages from R Bioconductor (107,108). For CpG level comparison, percentage of methylation of individual CpGs was calculated using the methylKit package (109) and the coverage files from the bismark aligner (110). To prevent PCR bias and increase the power of the statistical tests, we discarded bases with coverage less than 10 in all samples. The bisulfite conversion rate was calculated as the number of thymines (non-methylated cytosines) divided by coverage for each non-CpG cytosine, as implemented in methylKit. Differential methylation analysis of single CpGs between different groups was also performed in the same package and the CpGs with p-value less than 0.01 and delta beta value differences more than 25% defined as significant. In addition, density, PCA and correlation plots were also generated by R Bioconductor packages.

Differentially methylated regions were determined using DSS package (111–114) which outperforms other methods when sample size per group is small owing to the adoption of the Wald test with shrinkage for determining differentially methylated cytosines (115). We identified DMRs using the coverage files from bismark with a p-value threshold of 0.01 and delta beta value more than 25%. The individual CpGs and DMRs were annotated using Annotatr package (116).

For the enhancer analyses, differentially methylated regions (DMRs) between ALK tumor cells and Ctrl thymocytes were compared to ENCODE datasets for H3K4me1 and H3K27ac marks in murine thymus. We downloaded the ENCODE datasets from the ENCODE portal (72) (www.encodeproject.org) with the following identifiers: ENCFF666XCJ and ENCFF354DWX. Enhancers were predicted, and k-means clustering analysis were performed using deep Tools v2.0 (117). The ‘computeMatrix’command with the sub-command ‘scale-regions’ was used to generate the table underlying the heatmaps, using a 5-kb windows indicated by start (S) and end (E) of the DMR region. ‘plotHeatmap’ was used to visualize the table.

Region set enrichment analysis against publicly available datasets using the LOLA software was used to identify shared biologically patterns among differentially methylated regions (62). CpGs were merged into 1-kilobase tiling regions across the genome before LOLA analysis. The significant hyper- and hypomethylated regions obtained from our study were used as the query set and the set of all differentially methylated tilling regions were used as universe set. For a focused analysis, only region sets from embryonic stem cells, T lymphocytes and thymus in the LOLA core database were included. All the enrichments with a p-value below 0.05 were considered significant.

For network analysis, data were analyzed through the use of QIAGEN’s Ingenuity^®^ Pathway Analysis (IPA^®^, QIAGEN Redwood City, www.qiagen.com/ingenuity).

### Statistics

Data are represented as mean±SD, if not otherwise stated and were analyzed using GraphPad Prism (version 6, GraphPad Software, Inc. La Jolla, USA). To assess differences between groups, unpaired t-test or one- or two-way ANOVA followed by multiple comparison were used, depending on number of samples. Significance was defined according to following p-values: * p<0.05; ** p<0.01; *** p<0.001 **** p<0.0001. Not significant p-values are not shown. Survival statistics were analyzed using GraphPad Prism. Pairwise curve comparison to ALK tumors using log-rank calculations (Mantel-Cox test) were used to assess differences between the groups.

## Data availability

The RNA-seq and RRBS data sets will be deposited to the NCBI Gene Expression Omnibus (GEO) (118) and the accession numbers will be added to the final version.

## ACKNOWLEDGEMENT

This work was supported by funds of the Austrian Science Foundation (FWF) projects P27616 and I4066. SL is a fellow of the Postdoc Career Program at Vetmeduni Vienna.

## AUTHOR CONTRIBUTIONS

GE designed and supervised the study, E.R., P.H, M.R.H, S.L., T.D., L.H, M.Z., A.T., M.K., B.H.R performed experiments, R.S.T, C.H.D, G.S. performed bioinformatics analyses, G.T. provided the quantification of all imaging data, L.K.,C.S. provided mouse strains, C.B., C.S., W.E, supervised parts of the project, E.R,. G.S., G.E wrote the manuscript.

