## Supplementary material for "Requirement of DNMT1 to orchestrate epigenomic reprogramming during NPM-ALK driven T cell lymphomagenesis": Suppl_Fig legends

**­­** **Supplementary Figure 1. Immunophenotyping of ALK+ thymocytes and tumor cells. A,** FACS analysis showing CD4 and CD8 expression of thymocytes in 18-week old wildtype controls (Ctrl) versus ALK+ tumor-free (ALK tf) thymocytes and tumor cells (ALK tu) (gated for ALK expression) and their quantification of 3 biological replicates. Data are shown as mean±SD, ** p<0.01, **** p<0.0001 using two-way ANOVA. **B,** FACS analysis showing TCRβ and ALK expression of samples as in A, and their quantification of 3 biological replicates. Data are represented as mean±SD, ** p<0.01 using one-way ANOVA, n=3.

**Supplementary Figure 2. Immunophenotyping of peripheral ALK+ cells. A,** Percentage of ALK expression in all splenocytes as analyzed by nuclear FACS of splenocytes of 18-week old Ctrl, ALK tf and ALK tu samples. Representative histograms show expression levels of ALK in the three groups. Data are represented as mean±SD, ***p<0.001, ****p<0.0001 using one-way ANOVA, n=3.

**B,** FACS analysis of ALK and TCRβ (left panel) as well as CD4 and CD8 expression (middle panel) of splenocytes isolated from 18-week old tumor free mice (ALK tf) (upper panel) or ALK tumor mice (ALK tu) (lower panel) and the quantification of 3 biological replicates (right panel).

**Supplementary Figure 3. Different ALK tumor stages. A,** Thymus or tumor to body weight ratios in percent of Ctrl and tumor-free ALK (ALK tf) mice and of ALK tumor mice with small tumors (ALK sm) or end-term tumors (ALK tu). Graphs show mean±SD, *p<0.05. **p<0.01 using one-way ANOVA, followed by pair-wise comparison to Ctrl. **B,** Representative immunohistochemical ALK staining of Ctrl and ALK tumor stages as in **A,**.

**Supplementary Figure 4. Immunophenotyping of KO and ALKKO mice. A,** Percentage of ALK+ cells in 18-week old ALKKO compared to Ctrl and KO thymi. Histograms (right) indicate ALK expression levels in ALKKO compared to Ctrl and KO cells. Data are represented as mean±SD, ****p<0.0001 using one-way ANOVA, followed by unpaired t-test, n=3. **B,** FACS analysis showing representative CD4 and CD8 expression of thymocytes in 18-week old Ctrl, KO and ALKKO mice and the quantification of 3 replicates. Data are shown as mean±SD, two-way ANOVA. **C,** FACS analysis showing TCRβ and ALK expression of thymocytes isolated from Ctrl, KO and ALKKO thymi. The graphs show quantifications of 3 biological replicates. Data are represented as mean±SD, one-way ANOVA, no significant differences.

**Supplementary Figure 5. Differential gene expression. A,** Principal component analysis (PCA) of RNA-seq data of Ctrl, KO, ALKKO thymi and ALK tumors indicating the first two components with the largest part of variation in gene expression between the different genotypes. **B,** Venn diagrams showing unique and shared differentially expressed genes in different genotypes that are significantly upregulated (left) or downregulated (right) (FDR<0.05, absolute log_2_(FC)>1) compared to ALK tumors (upper panels) or compared to Ctrl (lower panels) samples. **C,** UpSet plot of intersections between deregulated genes of different pairwise comparisons. The bar chart on the left indicates the total number of deregulated genes between the pairwise comparisons of different genotypes. The upper bar chart indicates the number of intersected deregulated genes. The bar graph on the left depicts the number of deregulated genes per comparison. The red box highlights the number of genes commonly deregulated in ALK as well as ALKKO relative to Ctrl.

**Supplementary Figure 6. Changes in DNA methylation levels upon *Dnmt1* knockout. A,** Global DNA methylation analysis using dot blot and immunodetection with an antibody directed against 5mC (left) or methylene blue staining as DNA loading control (right). Hydroxymethylated (hmCpG) and methylated (mCpG) oligos were used as antibody control. 250 ng of DNA isolated from Ctrl, KO, ALK and ALKKO thymi and tumors were spotted onto the membrane. Analyses were performed in technical and biological triplicates. **B,** qRT-PCR of *IAP* expression in Ctrl, KO, ALK tumor and ALKKO samples. Analyses were performed in biological triplicates (n=3). Data are represented as mean±SD, *** p<0.001, **** p<0.0001, one-way ANOVA, followed by unpaired t-test. **C,D,** Annotation of hyper- (red) and hypo- (turquois) methylated CpG sites related to known gene annotations and CpG island characteristics for ALK versus Ctrl samples. **E,F,** Annotation of hyper- (red) and hypo- (turquois) methylated CpG sites related to known gene annotations and CpG island characteristics for ALKKO versus Ctrl samples.

**Supplementary Figure 7. Gating strategy for FACS analysis of thymocytes and tumor cells. A,** Representative gating strategy for the analysis of ALK negative thymocytes from Ctrl and KO mice. Cells were first gated for single cells before specific surface marker expression patterns were analyzed. **B,** Representative gating strategy for the analysis of ALK positive thymocytes and tumor cells from ALK and ALKKO mice. Ctrl samples were used as negative control for definition of ALK expressing cells. Expression of specific surface markers (CD4, CD8, CD25, CD44) was analyzed in ALK positive cells.

**Supplemental Table 1.** Gene list of significantly deregulated genes (FDR<0.05, absolute log_2_(FC)>1) between ALK tumors and Ctrl samples. The table shows the gene symbol, the ENSEMBLE gene ID, the mean of normalized counts of all samples (baseMean), log2 fold change, log fold change standard error (lfcSE), Wald statistics (stat), the p value and the false discovery rate-adjusted P value (FDR).

**Supplemental Table 2.** Gene list of significantly deregulated genes (FDR<0.05, absolute log_2_(FC)>1) between ALKKO and Ctrl samples as in Supplementary Table 1.

**Supplemental Table 3**. List of genes that showed significant deregulated gene expression (FDR<0.05, absolute log_2_(FC)>1) between ALK tumors and Ctrl thymocytes and that were located close to DMRs, which overlapped with active enhancers in thymi based on ENCODE ChIP-seq datasets for H3K4me1 and H3K27ac.
