## Supplementary figures and images for "Requirement of DNMT1 to orchestrate epigenomic reprogramming during NPM-ALK driven T cell lymphomagenesis"

### Suppl_Fig1_FACS.pdf

#### Supplementary Figure 1

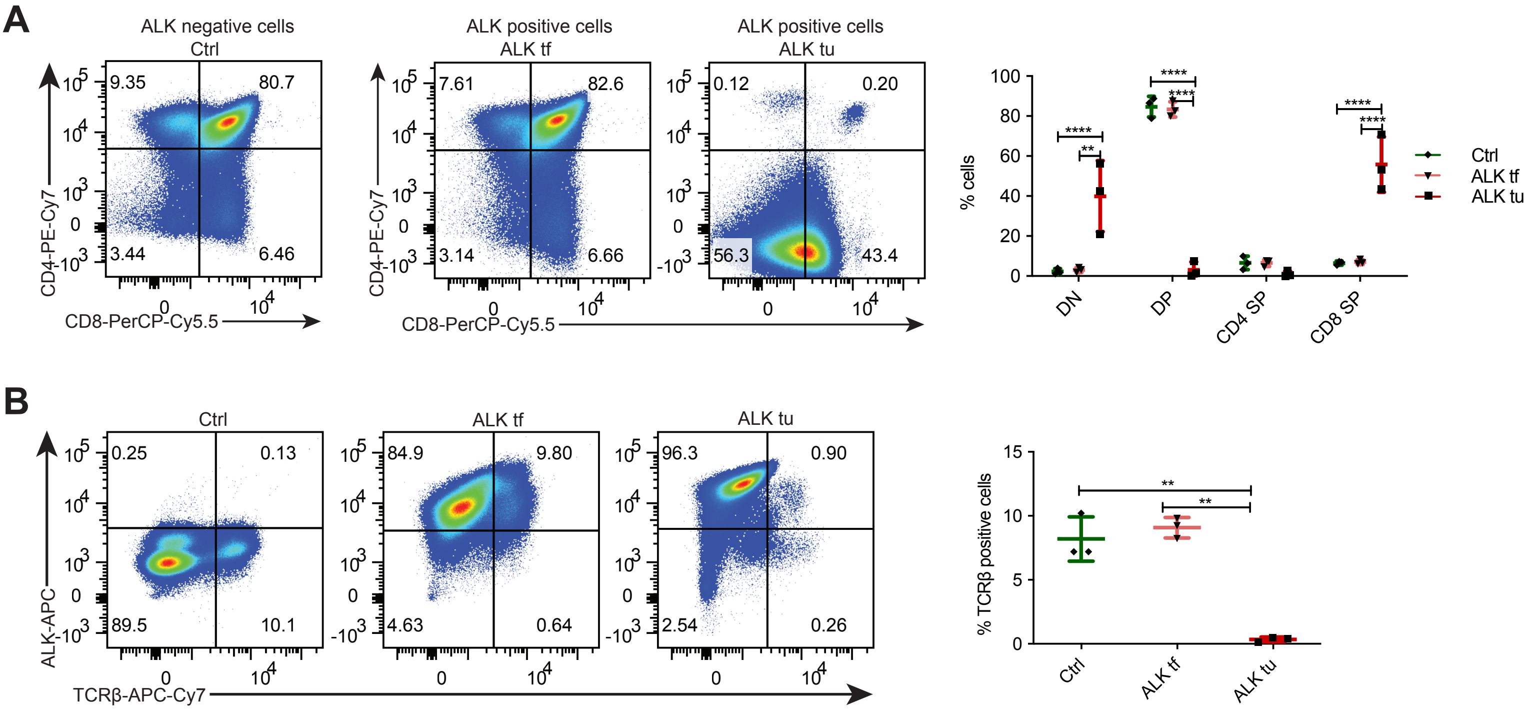

### Suppl_Fig2_Spleen_FACS.pdf

#### Supplementary Figure 2

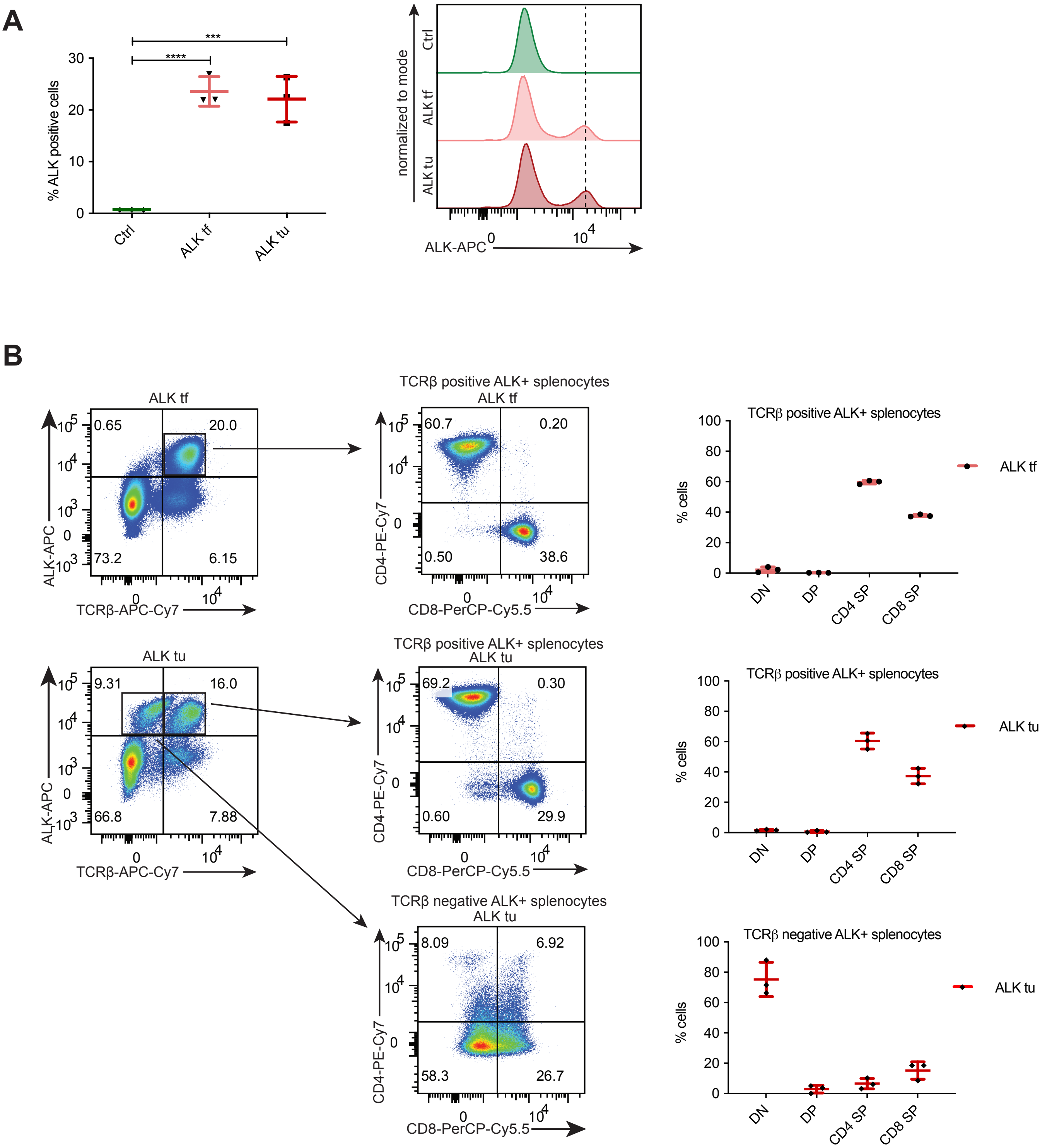

### Suppl_Fig3_opt.pdf

#### Supplementary Figure 3

A

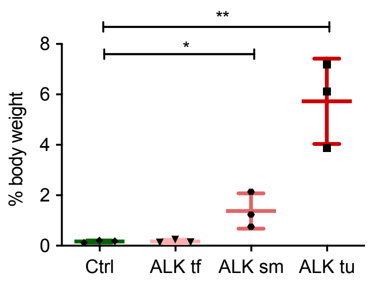


B

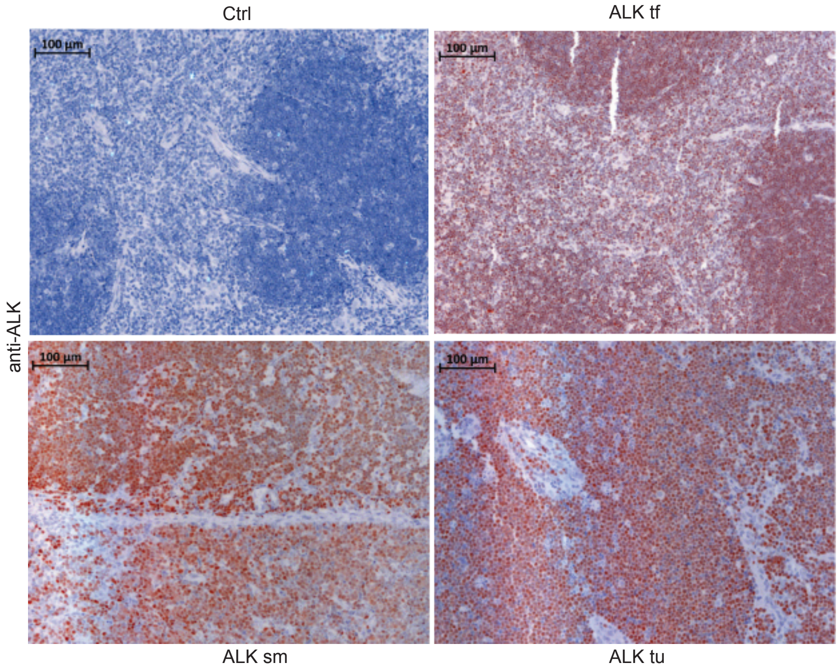

### Suppl_Fig4_FACS_mouse.pdf

#### Supplementary Figure 4

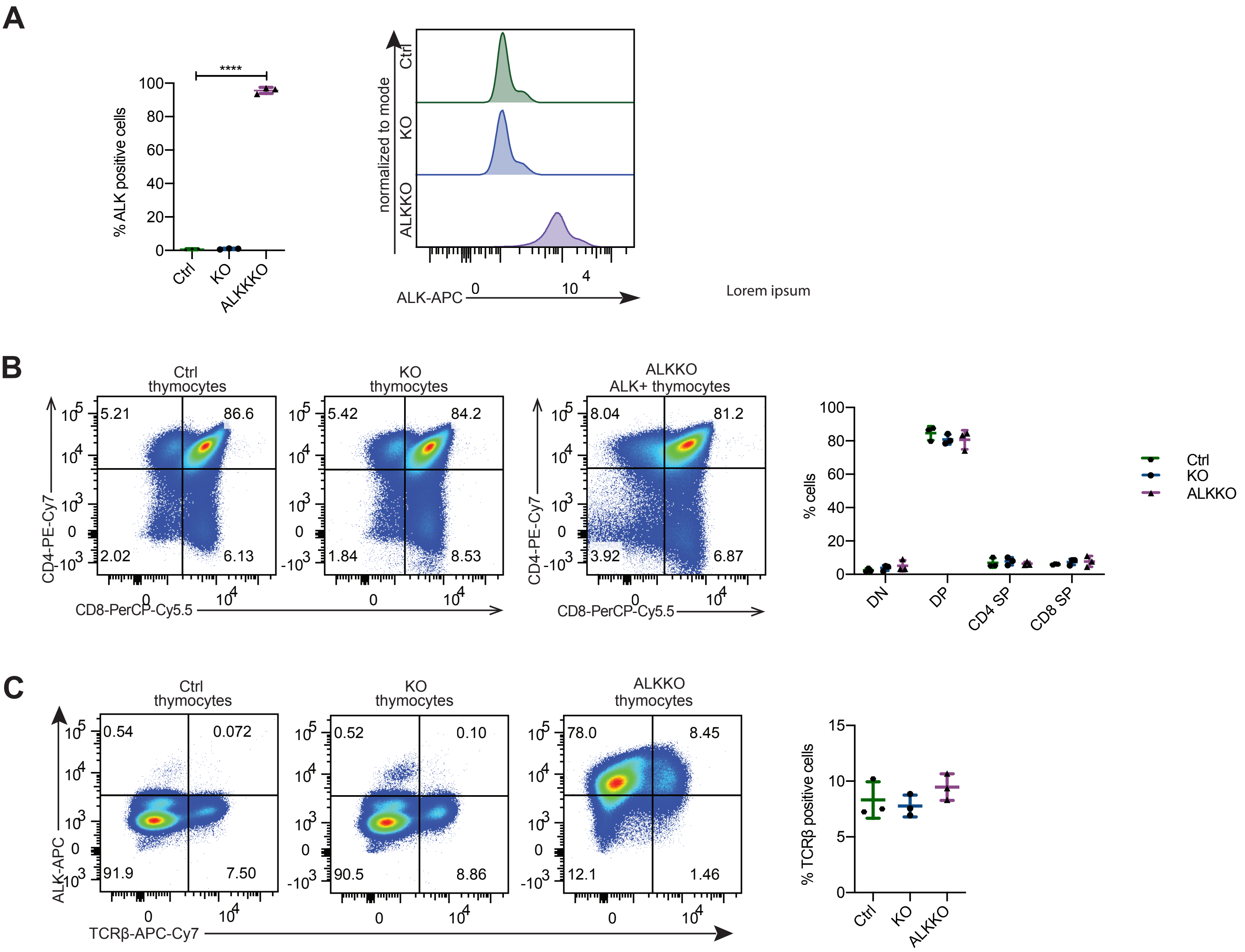

### Suppl_fig5_RNASeq.pdf

#### Supplementary Figure 5

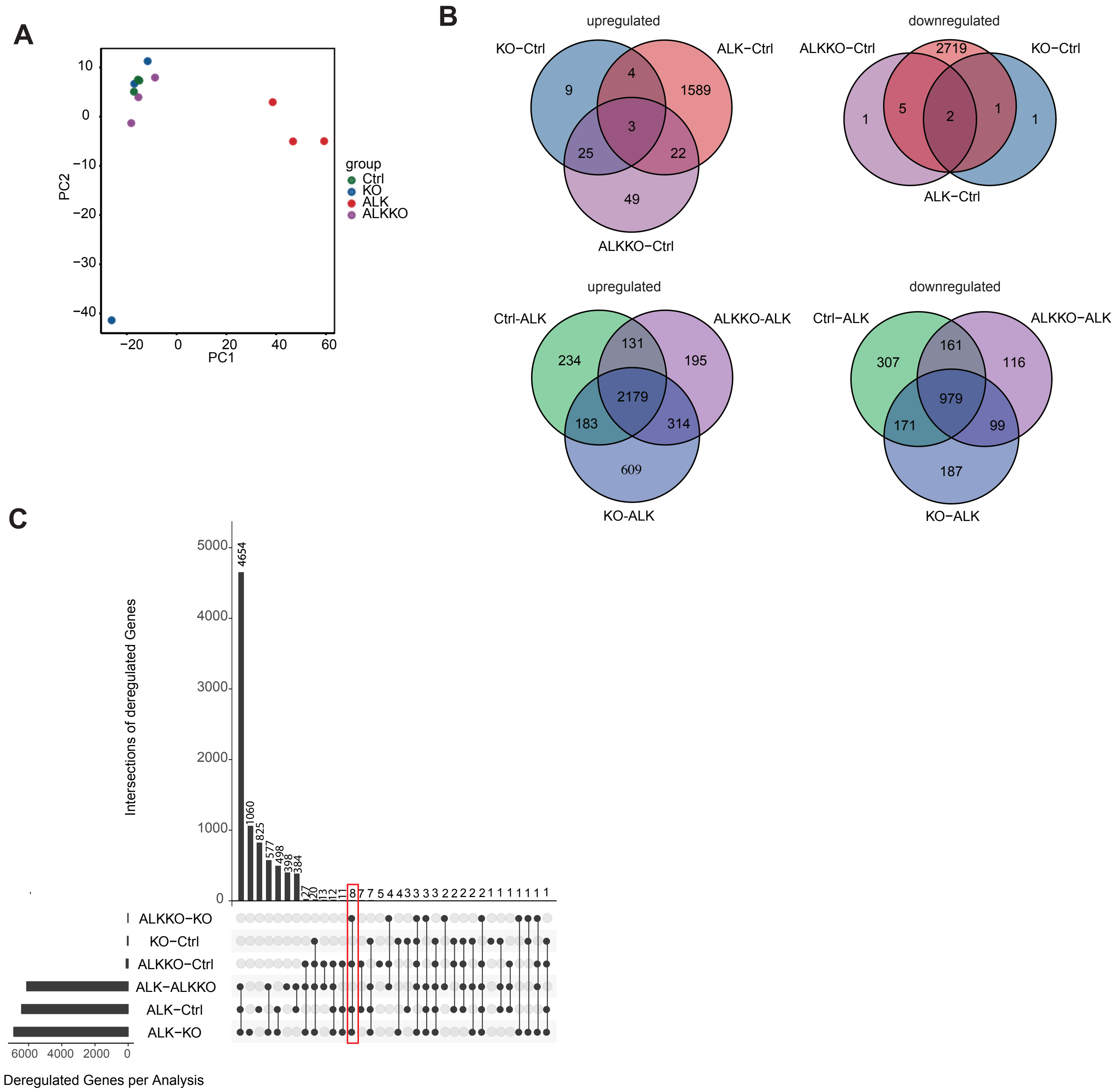

### Suppl_Fig6_methylation.pdf

#### Supplementary Figure 6

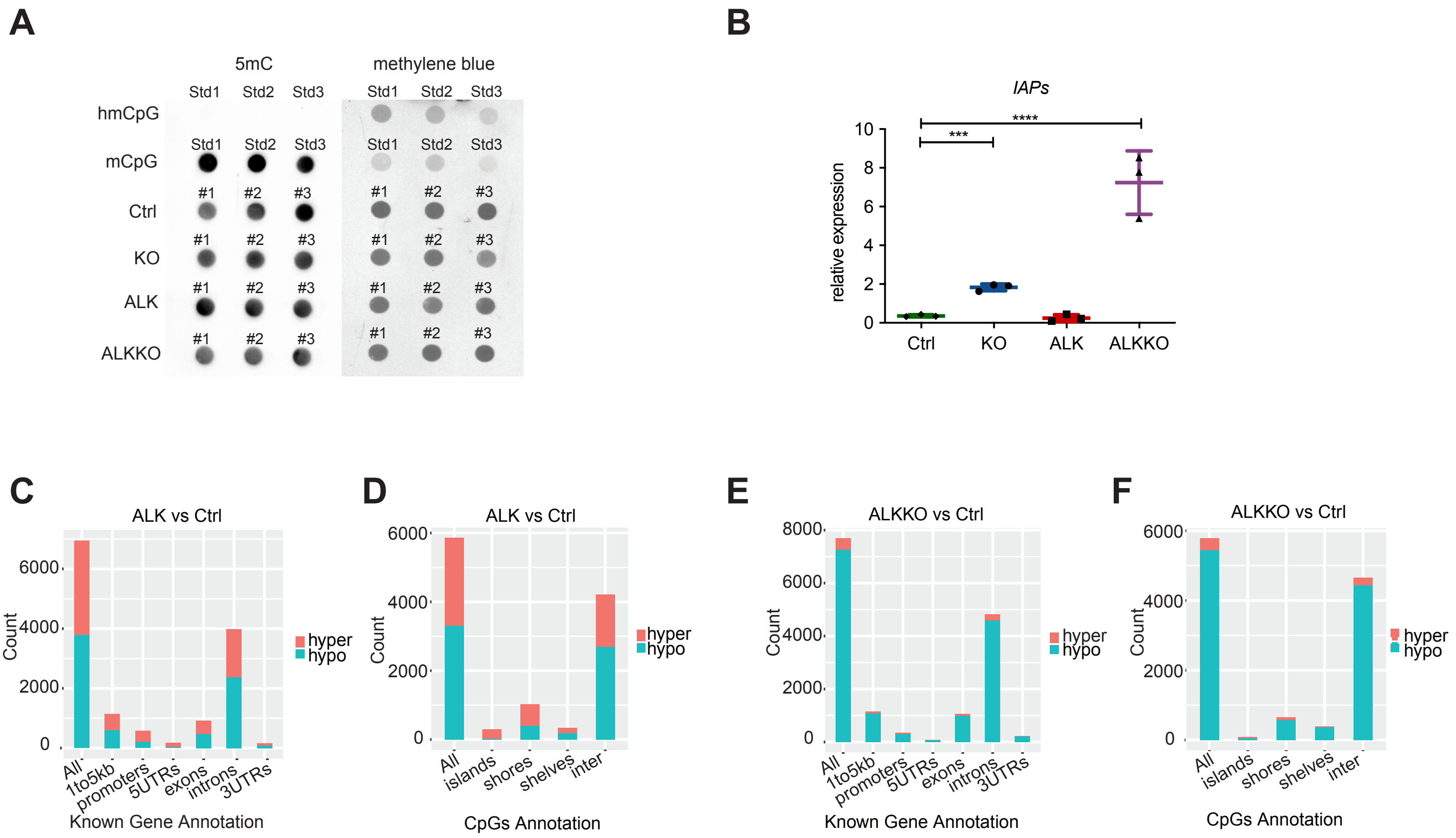

### Suppl_Fig7_gating_strategy.pdf

##### Supplementary Figure 7

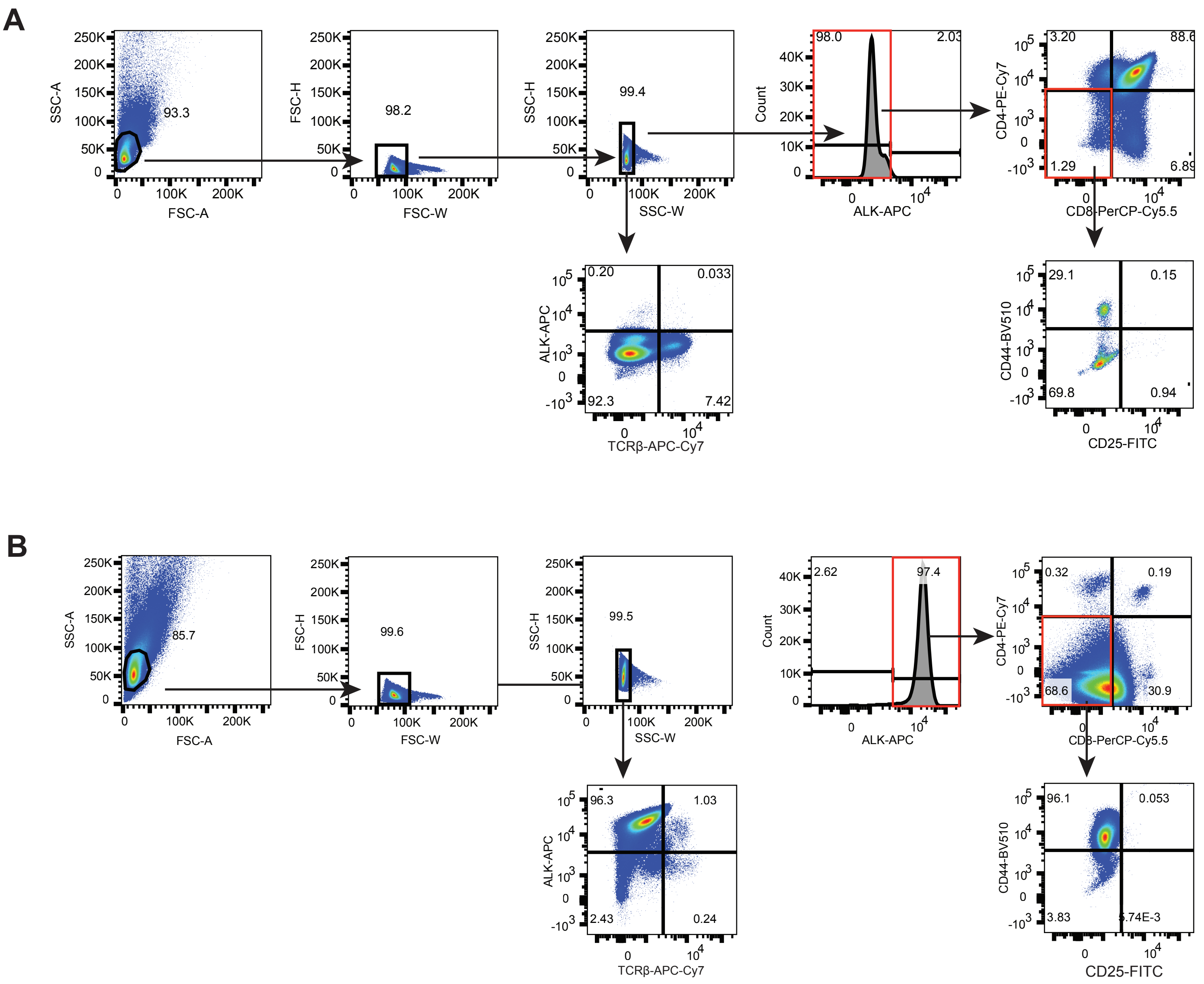
